# Immune–vascular dialogue coordinates muscle maintenance following damage in *Drosophila*

**DOI:** 10.1101/2025.06.25.661466

**Authors:** Nourhene Ammar, Olivier Josse, Aurélien Guillou, Emma Leroux, Kévin Macé, Jessica Perochon, Hadi Boukhatmi

## Abstract

In skeletal muscle, immune and vascular responses are essential for regeneration, yet the cellular and molecular mechanisms that coordinate their activities to restore tissue function remain unclear. Here, using *Drosophila*, we uncover a multi-organ signaling program that integrates muscle fibers, macrophages, vascular-like tracheal cells, and the extracellular matrix (ECM) to maintain muscle function in response to acute damage. Muscle injury induces rapid macrophage recruitment and secretion of the FGF-like ligand Branchless (Bnl), which activates FGF/FGFR signalling in tracheal cells and promotes their targeted expansion toward damaged muscle. Concomitantly, macrophages deposit ECM components on the damaged muscle, including Collagen IV, forming a localized microenvironment that restricts Bnl ligand diffusion and facilitates directed tracheal remodelling. Genetic disruption of macrophage-derived Bnl or macrophage-mediated ECM deposition abolishes tracheal remodelling and compromises muscle function following injury. Together, these findings reveal a novel immune–vascular communication program in which macrophages coordinate fibroblast growth factor signalling and ECM organization to shape muscle microenvironment and preserve muscle function following acute damage.

## Introduction

Effective tissue repair relies on precise communication and coordination across multiple cell types and tissue systems [1]. Skeletal muscle provides a prime example of this interdependent cellular network, as successful regeneration depends on tightly orchestrated interactions among diverse cell populations, including macrophages, muscle stem cells (MuSCs; also known as satellite cells), and the fibro/adipogenic progenitors (FAPs) [2, 3]. Following injury, infiltrating macrophages promote muscle regeneration by signalling to MuSCs and regulating their activation, proliferation, and differentiation [4, 5]. In parallel, extensive remodelling of both the vasculature and the extracellular matrix (ECM) is required to meet increased metabolic demands and provide structural support [6, 7]. While immune cells, vascular network, and ECM remodelling are all known contributors to muscle repair; how these systems are mechanistically integrated *in vivo* remains poorly understood. Addressing this question has been limited by the lack of experimental systems that enable to track and manipulate multiple interacting cell types *in vivo* in response to muscle damage.

*Drosophila* has emerged as a powerful *in vivo* model for dissecting multi-organ communication during regeneration, owing to its capacity to combine precise genetic manipulation with high-resolution imaging [8]. As in vertebrates, adult *Drosophila* indirect flight muscles (IFMs) harbor MuSCs that support muscle maintenance and have been implicated in regenerative responses following injury [9–12]. Studies, including our own, have shown that macrophage-like immune cells, termed hemocytes, populate the IFMs under homeostatic conditions [9, 13]. In addition to hemocytes, the IFMs are infiltrated by a sophisticated network of air-filled tracheal cells, which function analogously to the vertebrate vascular and respiratory systems by ensuring tissue oxygenation [14, 15]. While tracheal branching has been extensively characterized at the morphological, cellular, and genetic levels during development [16, 17], recent work has revealed that adult tracheal networks retain a remarkable capacity for dynamic remodelling in response to tissue damage, including during regeneration of the adult gut [18–20]. Finally, IFMs are ensheathed by a specialized basement membrane enriched in conserved ECM components, which provide structural integrity and mechanical support to the contractile apparatus [15, 21]. These features provide an *in vivo* context in which interactions between muscle fibres, immune cells, vascular-like trachea, and the ECM can be explored during the muscle injury response.

We show that muscle damage triggers rapid and sustained hemocyte recruitment, which is required for targeted tracheal remodelling through FGF/FGFR signalling, enabling selective recognition and support of damaged fibers. In parallel, hemocytes deposit ECM components around injured fibres, which spatially confine FGF-like ligand and promote tracheal recruitment. Disrupting tracheal remodelling or hemocyte-mediated ECM delivery compromises muscle performance after injury, revealing an immune–muscle–vascular signalling module that couples metabolic support to ECM organization to preserve muscle function upon damage.

## Results

### Elevated Mhc detection identifies damaged muscle fibers following injury

In order to assess the functional consequences of flight muscles injury over time, we conducted flight assays at 5- and 20-days post-injury (DPI), two widely separated time points (Figure S1 A). Flight capability was markedly reduced at 5 DPI compared with uninjured controls. However, no further decline was observed between 5 and 20 DPI (Figure S1 A), raising the possibility that active tissue responses could be engaged to stabilize muscle function following damage. To investigate these responses, we first sought to establish a framework to identify damaged muscle fibers and monitor the cellular changes associated with injury. In vertebrates, re-expression of embryonic Myosin Heavy Chain (MyHC-emb) in damaged fibers is a common marker of muscle injury and regeneration [22]. To assess whether Myosin Heavy Chain (Mhc) detection could similarly serve as an indicator of muscle damage in *Drosophila*, we analysed its expression following injury. Our results show that Mhc detection is increased in injured muscles compared with neighbouring intact fibers (Figure S1 B-C’). The elevated Mhc signal persists for at least 20 DPI (Figure S1 B-D’). To further evaluate the status of damaged muscles we examined the expression of the myogenic transcription factor (TF) Myocyte Enhancer Factor-2. In contrast to the increased Mhc signal, nuclear Mef2 expression is lost in damaged fibers by 5 DPI and remains undetectable at 20 DPI (Figure S1 B-D’’).

In *Drosophila*, several Mhc isoforms are generated via alternative splicing of a single *Mhc* gene [23]. Analysis of GFP-tagged Mhc isoforms revealed that only the shorter adult isoforms K, L and M displayed increased detection following muscle damage (Figure S1 E-G’’, H and I) [23, 24]. Furthermore, a similar enhanced Mhc signal was observed in detached IFMs of Naa30A mutants, a genetic model of muscle detachment (Figure S1 J-M’’) [25]. Thus, elevated Mhc detection provides a sensitive marker of muscle fiber damage in adult IFMs.

We next asked whether the injured muscle fibers remain alive following damage. To address this question, we examined both nascent transcription and apoptotic activity within injured muscles (Figure S2). Active *mef2* transcription sites were detected in 35% of nuclei within injured muscles at 5 DPI, compared with 75% of nuclei in neighbouring intact fibers (Figure S2 A-B). In parallel, we found no detectable signal for the apoptotic marker Drosophila caspase-1 (Dcp1) at this stage. In contrast, elevated Dcp1 staining was observed in 28% of damaged fibers at 20 DPI (Figure S2 C-G). Together, these observations indicate that injured fibers remain viable during the early phase following injury, providing a temporal window in which cellular responses to muscle damage can be investigated.

### Hemocytes are recruited to the injured adult flight muscle

The hemocytes are among the cell populations associated with adult IFMs [9, 13], however their contribution in IFMs injury response remains largely unexplored. To address this, we started by tracking these cells at various time points after injury using the hemocytes reporters Peroxidasin (*Pxn*) (Figure 1 A-C’’) and Hemolectin (*Hml*) (Figure S3 A-B’’). Co-staining for these markers and Mhc revealed extensive recruitment of hemocytes to the injured muscle fibers (Figure 1 A-C’’; Figure S3 A-B’’). To further characterize the hemocytes recruitment dynamics, we examined the Zinc finger homeodomain transcription factor Zfh1 reporter (*Enh3-Gal4 > UAS-mCD8GFP*), which also labels hemocytes [9, 26]. Enh3+ cells were similarly recruited to damaged fibers (Figure 1 D-F’’). We then compared hemocytes recruitment kinetics based on the expression of these three markers (Figure 1 G). Enh3+ hemocytes displayed recruitment kinetics similar to those of Pxn+ cells, consistent with our observation that all Pxn+ cells are Enh3+ (Figure 1 G and Figure S3 C-D’’). In contrast, although Hml+ hemocytes were initially recruited with similar kinetics to Pxn+ and Enh3+ cells, their abundance significantly declined by 5 DPI (Figure 1 G). Accordingly, only 55% of Pxn+ hemocytes remained Hml+ at this stage (Figure 1 G’ and Figure S3 A-D’’). Together, these results reveal that the hemocyte population recruited to injured muscles is heterogeneous and comprises cells expressing distinct molecular markers with different temporal dynamics.

**Figure 1.**
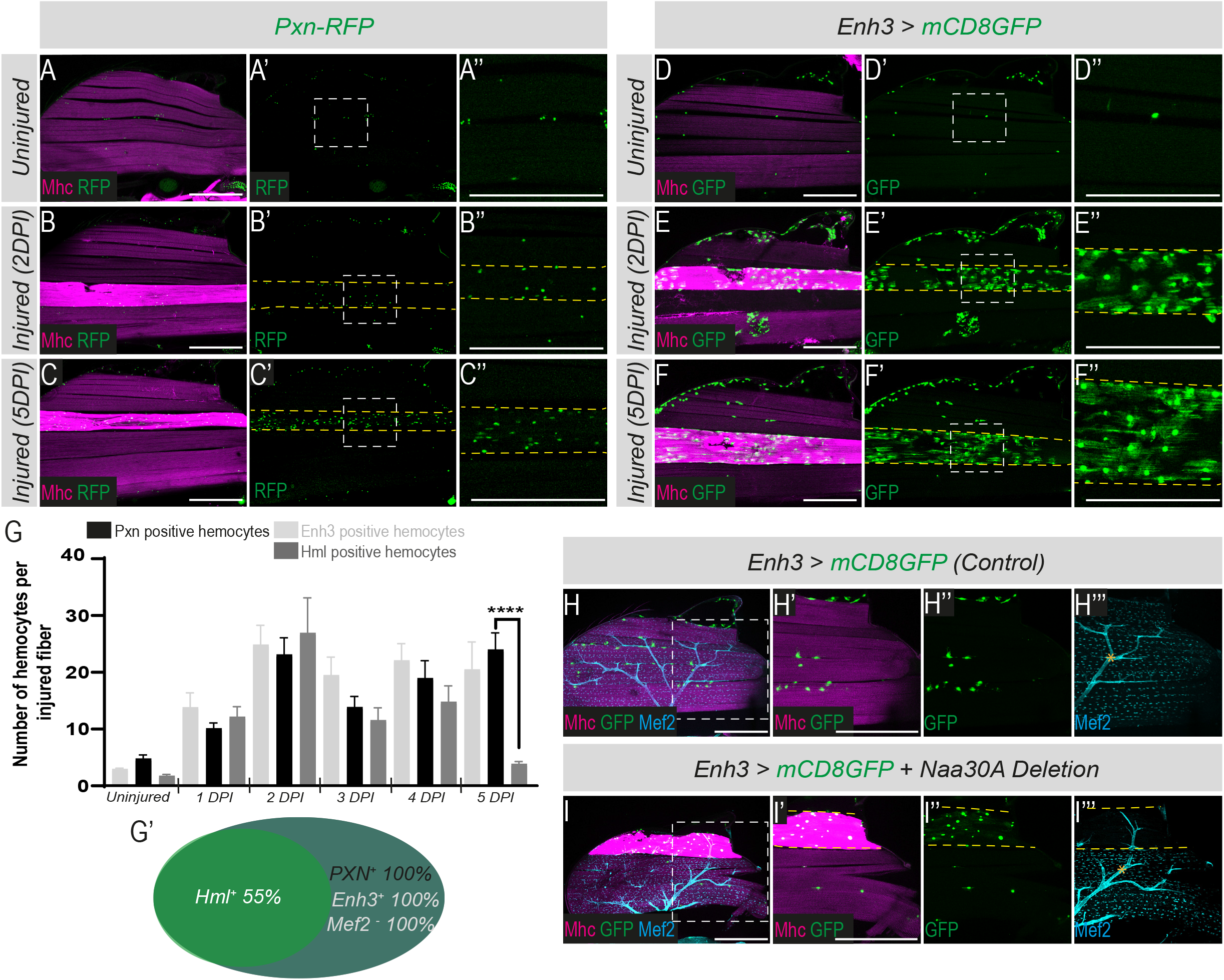
Hemocytes are specifically recruited to the damaged muscle fiber. **(A-F).** Adult indirect flight muscles (IFMs) imaged under uninjured conditions and at 2-and 5-days post-injury (2DPI and 5DPI). Mhc (magenta) marks muscles. Hemocytes (green) are visualized using the following reporters: *Pxn-RFP* in (A-C’’) and *Enh3-Gal4; UAS-mCD8GFP* in (D-F’’). Dashed yellow outlines indicate injured muscle fibers showing elevated Mhc detection. Boxed regions are magnified 5X. **(G).** Time-course analysis of hemocytes recruitment at 1, 2, 3, 4 and 5 DPI reveals heterogeneous kinetics of response to muscle damage. Enh3-positive, Pxn-positive, and Hml-positive hemocytes are represented by light gray, black, and dark gray bars, respectively (****p < 0.0001, Student’s t-test; n ≥ 10 heminota per condition). **(G’).** Schematic representation of the proportion of recruited hemocytes expressing different reporters at 5DPI (n ≥ 10 heminota per condition from three independent biological replicates). None of Enh3+ cells express Mef2. Quantification was performed on more than 400 cells from three independent biological replicates. **(H-I’’’).** Distribution of hemocytes visualized with *Enh3-Gal4; UAS-mCD8GFP* (green) in control muscles (H-H’’’) and *Naa30A* mutants (I-I’’’). Muscles and muscle nuclei are stained with Mhc (magenta) and Mef2 (cyan) respectively. Scale bars: 200μm. Asterisks in panels H’’’ and I’’’ mark non-specific signal detected in motoneurons.

We previously showed that Zfh1/Enh3-Gal4 labels both MuSCs and hemocytes within the IFMs [9]. We therefore asked whether any of the recruited Enh3+ cells correspond to MuSCs. None of the recruited Enh3+ cells detected at injured fibers express Mef2, indicating that they do not belong to the muscle lineage (Figure S3 E-F’’). To further define their identity, we used the lineage-tracing system ReDDM (Repressible Dual Differential Stability Markers) [28], which relies on the differential stability of a short-lived membrane-bound GFP (mCD8-GFP) and a long-lived nuclear RFP (H2B-RFP) to trace cell lineage and progeny production (Figure S3 G-I’) [27]. All recruited Enh3+ cells remained GFP+/RFP+ following injury (5 DPI), indicating that they do not generate newly differentiated progeny through asymmetric division (Figure S3 H-I’). Consistently, none of the recruited Enh3+ cells expressed the mitotic marker phospho-Histone H3 (PH3) following injury (Figure S3 J-K’). These data show that the recruited Enh3+ cells correspond to hemocytes rather than activated MuSCs.

To determine whether hemocyte recruitment is a general feature of muscle damage and not restricted to the mechanical stab injury paradigm, we also analysed their comportments in the *Naa30* mutants [25]. We observed a selective recruitment of Enh3+ hemocytes to damaged muscle fibers identified by strong Mhc staining and loss of nuclear Mef2 expression (Figure 1 H-I’’’). These findings indicate that hemocyte recruitment is a common response to muscle damage and is not limited to stab induced injury.

### Hemocytes recruitment coordinates with tracheal remodelling following muscle injury

We observed that hemocytes are closely associated with muscle tracheal cells under homeostatic conditions (Figure 2 A-A’). Previous studies have shown that adult tracheal cells can undergo dynamic remodelling and extend new branches toward damaged regions to support gut regeneration [18–20]. We thus asked if tracheal remodelling also occurs in response to IFMs injury and whether this process involves any dialogue between tracheal cells and hemocytes. Our analysis revealed that muscle damage, whether caused by physical injury or in *Naa30A* mutant animals, leads to a significant increase in tracheal coverage of the injured fiber (Figure 2 B-D and Figure S4 A-B’). The invasion is directed toward the injury site, with the highest branching density close to the damaged area and progressively decreasing with distance from the injury (Figure 2 E). We further observed that hemocytes remain spatially close to the tracheal branches, even under injured conditions, suggesting sustained interactions between these two cell types during the muscle damage response (Figure 2 C’’).

**Figure 2.**
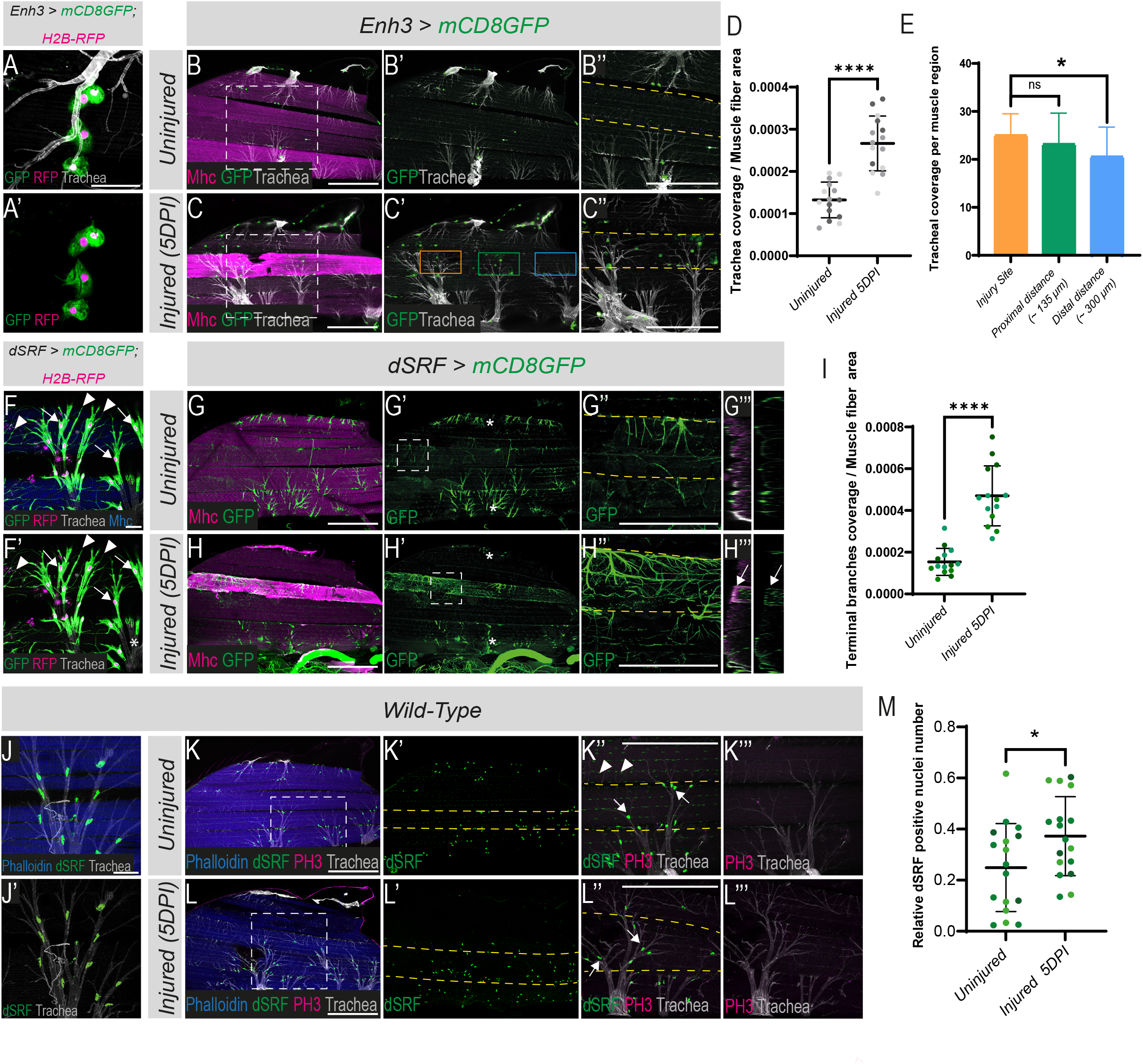
Muscle damage induces directed tracheal remodelling. **(A-A’).** Hemocytes (green/magenta; *Enh3-Gal4, UAS-mCD8GFP; UAS-H2B-RFP*) are associated to the muscle tracheal cells (gray, autofluorescence). Scale bars: 25 μm. **(B-C’’).** Distribution of tracheal branches (gray, autofluorescence) and hemocytes (green, *Enh3-Gal4; UAS-mCD8GFP*) in uninjured IFMs (B-B’’) and at 5DPI (C-C’’). Muscles are labeled with Mhc (magenta). (B’’, C’’). Boxed regions from (B and C), magnified 2X. Scale bars: 200 μm. **(D).** Quantification of tracheal coverage in (B) and (C) (****p < 0.0001, Student’s t-test; n = 15 heminota per condition; light, dark, and intermediate shading indicate data points from three independent replicates). **(E).** Quantification of tracheal coverage in (C) at the injury site (orange box), adjacent to the injury site (green box, approximately 135 μm away from the injury site), and at distant regions (blue box, within 300 μm of the lesion border) (*p= 0.0142, Student’s t-test; n=19 heminota). **(F-F’).** dSRF-Gal4 marks muscle-associated terminal tracheal cells (TTCs). TTC membranes (green) and nuclei (magenta) are indicated by arrowheads and arrows, respectively (*dSRF-Gal4, UAS-mCD8GFP; UAS-H2B-RFP*). dSRF-Gal4 is inactive in main tracheal branches (asterisks in F’). Muscles are labeled with Mhc (blue). Scale bars: 25 μm. **(G-H’’’).** TTCs distribution in IFMs (green, *dSRF-Gal4, UAS-mCD8GFP*) in uninjured muscles (G-G’’’) and at 5DPI (H-H’’’). Muscles are labeled with Mhc (magenta). (G’’, H’’). 4.5X magnifications of boxed regions in (G’) and (H’). Scale bar: 200 μm. (G’’’, H’’’). Orthogonal views of (G’’) and (H’’). Arrow in (H’’’) indicates TTC enrichment at the injured muscle. Asterisks in panels G’ and H’ mark the fibers #1 and #5. **(I).** Quantification of TTCs coverage from (G) and (H) (****p < 0.0001, Student’s t-test; n = 14 heminota per condition; light, dark, and intermediate shading points represent data points from three independent replicates). **(J-J’).** dSRF (green) marks TTCs nuclei associated with IFMs. Scale bars: 25 μm. **(K-L’’’).** Distribution of TTCs nuclei (green) in IFMs under uninjured conditions (K-K’’’) and at 5DPI (L-L’’’). Muscles are labeled with phalloidin (blue); tracheae are visualized by autofluorescence (gray). Anti-PH3 (magenta) is used to assess cell division. (K’’, L’’). 2X magnification of boxed regions in (K) and (L). Scale bars: 200 μm. Arrowheads and arrows indicate dSRF expression in the muscle and TTCs nuclei, respectively. **(M).** Proportion of TTCs nuclei located within a selected muscle relative to the total number of TTCs nuclei in the imaged sample (*p = 0.036, Student’s t-test; n = 17 heminota per condition; light, dark, and intermediate shading indicate data points from three independent replicates).

To further investigate the tracheal response to injury, we focused on terminal tracheal cells (TTCs), specialized branches of the tracheal system known for their structural plasticity and ability to invade tissues, including muscle fibers, to deliver oxygen [14, 15, 28]. TTCs can be reliably identified by their distinctive morphology and the expression of the transcription factor Serum Response Factor (dSRF) (Figure 2 F-F’ and J-J’). We visualized TTCs by expressing membrane-bound GFP under the control of dSRF-Gal4 (*dSRF-Gal4 > UAS-mCD8GFP*) (Figure 2 G-H’’’). We observed a high density of TTCs within injured muscle fibers, with the highest concentration near the injury site (Figure 2 H-H’’’ and I). Orthogonal views showed that the recruited trachea are found not only on the muscle surface but also extend branches into the injured fibers (Figure 2 G’’’-H’’’), consistent with the atypical muscle-tracheal organization optimized to deliver oxygen deep within the muscle, directly to sites of high metabolic demand [14, 15]. Labelling TTCs nuclei with anti-dSRF confirmed the significant enrichment of dSRF+ nuclei along the injured fibers comparing to the uninjured muscles (Figure 2 K-L’’ and M). The tracheal enrichment at the injured muscle results from TTCs branching and recruitment rather than proliferation, as no PH3+ cells were detected in the muscle trachea cells (Figure 2 K’’’-L’’’). Altogether, these data indicate that tracheal cells respond to muscle injury and target specifically the damaged fiber.

The tracheal enrichment at the injured muscles persists for at least 20 DPI (Figure S4 C-K’ and N). Recruited hemocytes first accumulated near the injury site and subsequently expanded along the damaged fiber (Figure S4 C-K). A comparison of hemocyte recruitment kinetics with tracheal coverage density revealed that hemocyte recruitment precedes tracheal coverage of the injured fibers (Figure S4 M and N). These data raise the possibility of an interplay between the two-cell population in supporting the injured muscle fiber.

### Tracheal invasion of injured muscle fibers is triggered by local hypoxia

The tracheal recruitment to injured muscle fibers (Figure 2) suggests a critical oxygen demand of the damaged muscle. If this is the case, local hypoxia within injured fibers could serve as a key cue for tracheal attraction and directional growth toward the site of injury, as seen in other physiological contexts [29]. To assess oxygen variations within the injured muscle, we first used the Synthetic HRE (Hypoxia response element) nuclear sensor (SyHREns). A hypoxia GFP reporter that serves as a proxy for evaluating intracellular oxygen fluctuations [30]. In normal condition, the SyHREns activity was primarily detected in TTCs, identifiable by both their triangular shape and expression of dSRF (Figure 3 A-A’’). Following injury, a pronounced enrichment of SyHREns-positive cells is detected surrounding injured muscle fibers, in close proximity to the recruited tracheal branches (Figure 3 B-D’’). These SyHREns+ cells corresponded predominantly to TTCs, although a small subset of recruited hemocytes also express the SyHREns reporter (Figure S5 A-B’’’). In contrast, the SyHREns reporter is not detected in the nuclei of damaged muscle fibers (Figure S5 C-D’). The high number of SyHREns+ cells around the injured fibers remained stable for at least 20 DPI (Figure 3 E), indicating that local hypoxia persists in parallel with the sustained tracheal remodelling observed over the same period.

**Figure 3.**
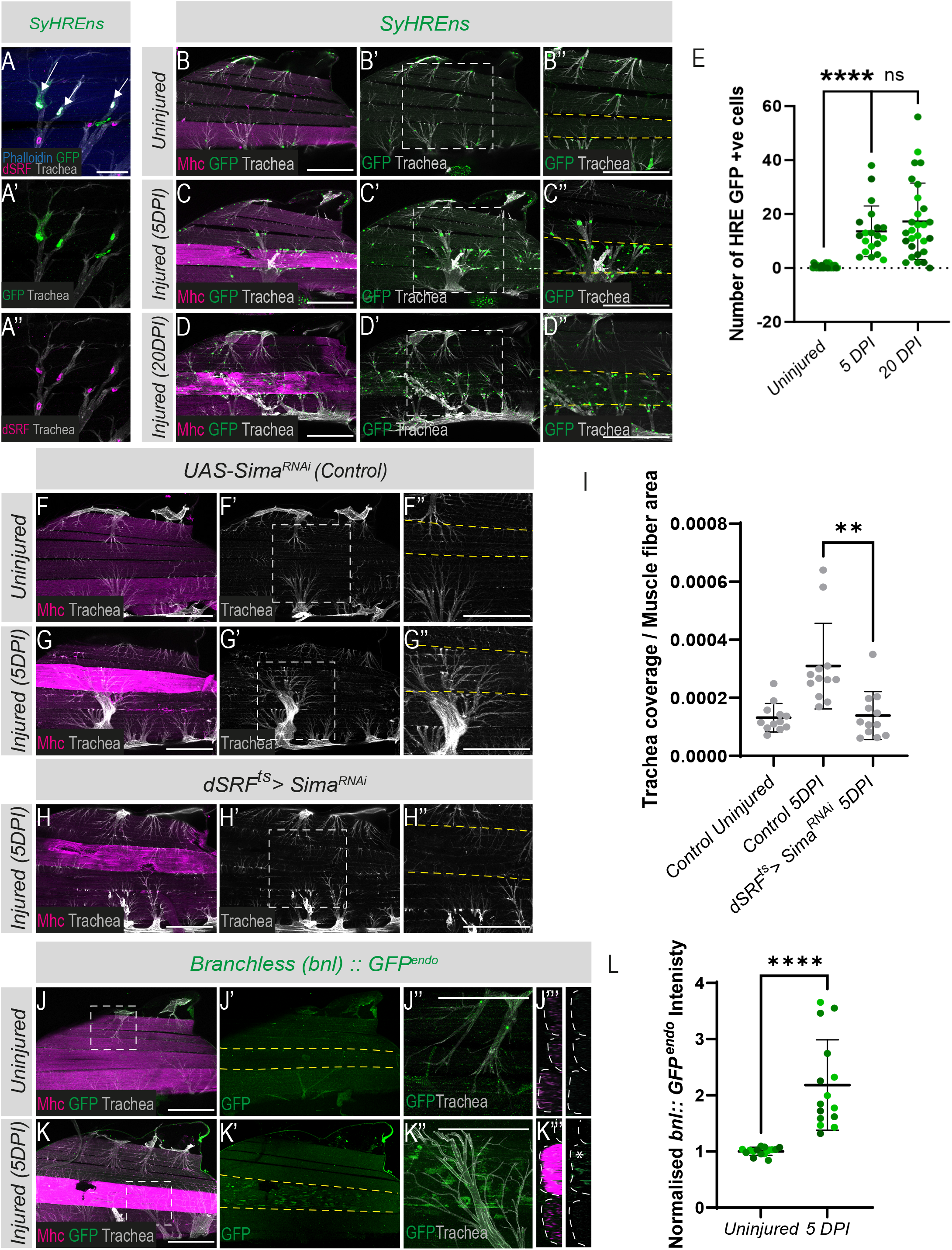
Local hypoxia in response to muscle damage drives tracheal branching. **(A-A’’).** *SyHREns* reporter activity (green) is detected in a subset of TTCs marked by dSRF (magenta, arrows). Scale bars: 25 μm. **(B-D’’).** Expression of the *SyHREns* reporter (green) in unijured IFMs (B-B’’), at 5DPI (C-C’’) and 20DPI (D-D’’). Muscles are labeled with Mhc (magenta), and tracheal branches are visualized via autofluorescence (gray). (B’’, C’’ and D’’). 2X magnification of the boxed regions in B’, C’ and D’. Scale bars: 200 μm. **(E).** Quantification of SyHREns positive cells (green) per injured muscle fiber (****p < 0.0001, Student’s t-test; n > 20 heminota per condition; light, dark and intermediate shading indicate data points from three independent replicates). **(F-H’’).** *sima* (HIF-1α) downregulation in TTCs (*dSRF-Gal4; UAS-Sima^RNAi^*) prevents tracheal recruitment at the injured fiber. (F’’, G’’ and H’’). 2X magnification of the boxed regions in E’, F’ and G’. Scale bars: 200 μm. **(I).** Quantification of trachea coverage (****p < 0.0001, Student’s t-test; n = 13 heminota per condition). **(J-K’’).** Detection of Bnl (green, *Bnl::GFP^endo^*) in control IFMs (J-J’’) and at 5DPI (K-K’’). Muscles are labeled with Mhc (magenta), and tracheal cells are visualized via autofluorescence (gray). (J’’, K’’). 4.5X magnification of the boxed regions in J and K. Scale bars: 200 μm. (J’’’, K’’’). Orthogonal views of (J) and (K). Asterisk in K’’’ indicates Bnl::GFP^endo^ enrichment at the injured muscle. **(L).** Quantification of normalized Bnl::GFP^endo^ intensity level in uninjured and injured muscle fiber at 5DPI (***p<0.0001, Student t-test, n = 15 heminota per condition from three independent replicates).

To determine the functional contribution of hypoxia signaling in these cell populations, we next down regulated the transcription factor Similar (Sima) -the *Drosophila* homolog of hypoxia-inducible factor-1α (HIF-1α)-a central regulator of hypoxic and angiogenic responses [29]. Silencing *sima* in TTCs strongly impaired tracheal expansion toward injured fibers but did not affect hemocyte recruitment to the damaged muscle (Figure 3 F-I and Figure S5 E-G). In contrast, depletion of *sima* in hemocytes neither affected hemocyte recruitment nor altered tracheal enrichment at the injury site (Figure S5 H-L). Together, these data indicate that activation of the hypoxia pathway in TTCs is the major driver of tracheal remodelling in response to injury.

Bnl is a well-established downstream effector of hypoxic signalling, deposited locally on hypoxic tissues to promote tracheal attraction [29]. We therefore hypothesised that if the injured muscle undergoes hypoxia, Bnl should accumulate locally at the damaged fiber to facilitate tracheal remodelling. Analysing the expression of an endogenously tagged *Bnl::GFP^endo^* allele [31] revealed marked enrichment of Bnl at injured muscle fibers compared to adjacent intact ones (Figure 3 J-K’’’ and L). Altogether, these data suggest that muscle damage induces local muscle oxygen starvation, which in turn drives tracheal cell recruitment to the injured tissue.

### Hemocytes deposit FGF-like ligand Bnl at injured muscles to mediate tracheal recruitment

During both embryonic and post-embryonic stages, tracheal guidance is regulated in a dose-dependent manner by local Bnl concentrations [29]. In adult flight muscle development, muscle-tracheal connections are established through the action of Bnl, which is secreted by muscle fibers and acts in a paracrine manner by binding to its receptor, FGFR/Breathless (Btl), expressed on tracheal cells [14]. The observed enrichment of Bnl in injured muscles (Figure 3 J-L) raises two key questions: Is Bnl/Btl signaling functionally involved in directing tracheal recruitment to damaged fibers? If so, what is the source of Bnl enrichment on the injured fibers? To begin addressing these questions, we conditionally overexpressed Bnl in a restricted subset of adult muscle fibers using the flip-out system combined with a regionally expressed driver (Figure S6 A-C and Materials and Methods section). This targeted overexpression was sufficient to recruit tracheal cells to the ectopic Bnl-expressing regions (Figure 4 A-C’), and the extent of this recruitment depended on the duration of Bnl expression: prolonged expression (5 days; Figure 4 C-C’) triggered more extensive tracheal remodelling than short-term expression (24 hours; Figure 4 B-B’). These findings showed that the adult tracheal system remains responsive to elevated Bnl levels in the IFMs, supporting the hypothesis that Bnl/Btl signalling directs tracheal pathfinding in response to muscle injury.

**Figure 4.**
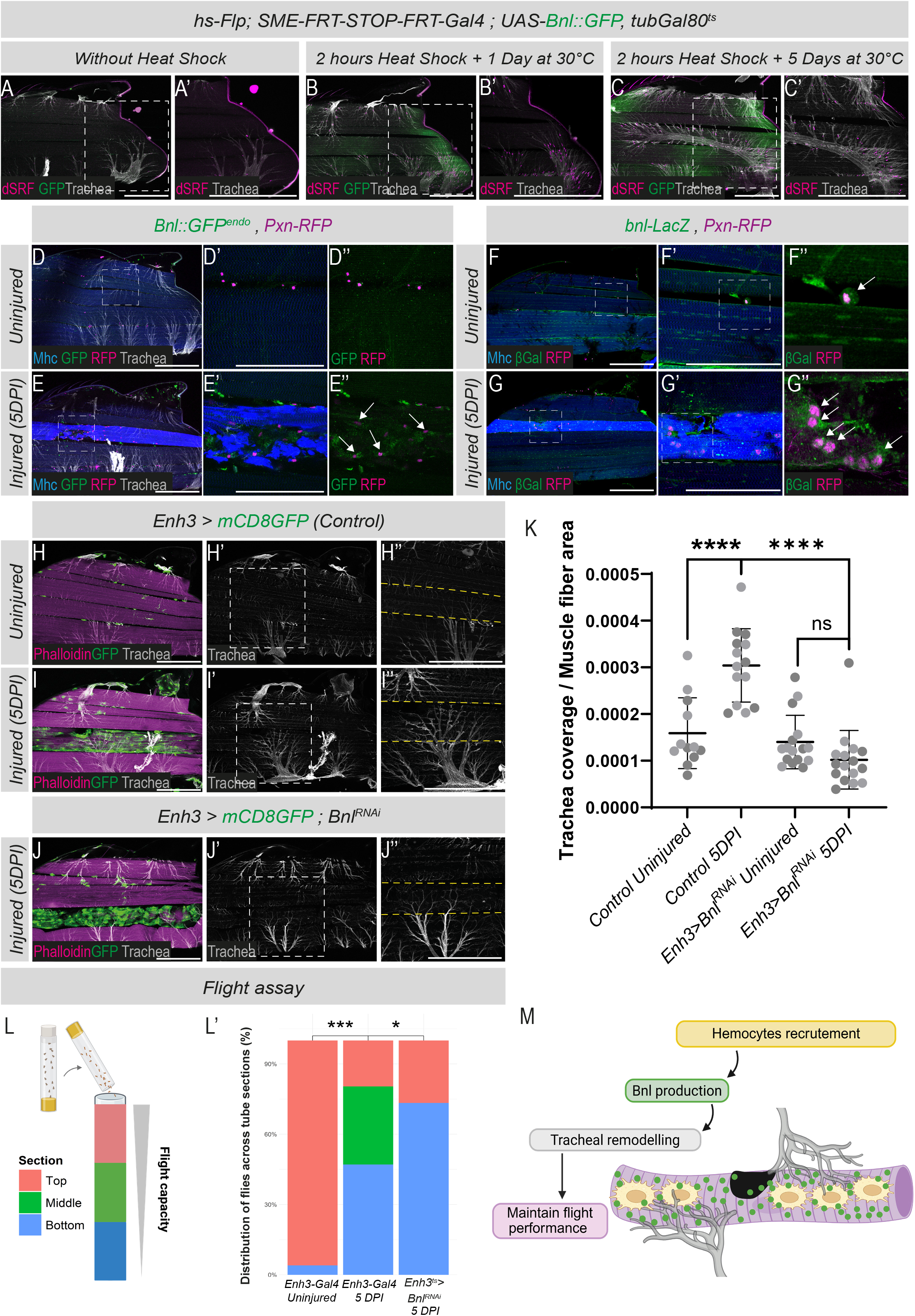
Hemocytes deliver Bnl to injured muscle to promote tracheal recruitment. **(A-C’).** Local overexpression of UAS-Bnl::GFP (green) in adult IFMs using *SME-Gal4* driver induces enrichment of terminal tracheal cells (TTCs; dSRF, magenta) and increases tracheal branching (gray), compared with controls (A). Scale bars: 200 μm. **(D-E’’).** Endogenous Bnl expression (green, *Bnl::GFP^endo^*) in uninjured IFMs (D-D’’) and at 5DPI (E-E’’). Muscles are labeled with Mhc (blue), tracheal branches are visualized via autofluorescence (gray), and hemocytes detected with the *Pxn-RFP* reporter (magenta). Arrows in E’’ indicate Bnl localization around hemocytes recruited to the injured muscle. Scale bars: 200 μm. **(F-G’’).** Expression of the *bnl-LacZ* reporter (green) is detected in IFMs associated hemocytes labelled with *Pxn-RFP* (magenta; arrows) in uninjured condition (F-F’’) and at 5DPI. **(H-J’’).** Downregulating *bnl* in the hemocytes (green, *Enh3-Gal4; UAS-mCD8GFP, UAS-Bnl^RNAi^*) do not alter hemocytes recruitment but decreases tracheal invasion to the damaged fiber as shown in (J’-J’’) and quantified in **(K)** (**** p< 0.0001, Student t-test; n ≥ 14 heminota from two independent replicates). Scale bars: 200 μm. **(L).** Schematic representation of the flight-assay setup used to assess flight performance. Flies are released at the top of a graduated cylinder, and landing position across three vertical zones is recorded. **(L’).** Quantification of the proportion of flies distributed across three sections under three conditions: *Control Uninjured*, *Control Injured (5DPI)* and *Enh3-Gal4; UAS-Bnl^RNAi^ Injured (5DPI)*. Bars indicate the percentage of flies landing in each of the three flight-assay zones (***p < 0.001, *p < 0.05, Chi-square test; n > 60 flies per condition per replicate. Data were obtained from two independent biological replicates). **(M).** Schematic model of hemocyte-driven tracheal recruitment required to maintain flight capacity following muscle injury.

We next sought to identify the cellular source responsible for Bnl accumulation in injured muscle fibers. Under wild-type conditions, Bnl::GFP^endo^ signal was detected in hemocytes, tracheal cells, and IFMs (Figure S6 D-D’). However, since Bnl is a diffusible protein, these observations alone cannot distinguish between cells that produce Bnl and those that merely take it up. To address this, we used the *bnl-lacZ* transcriptional reporter, which revealed active *bnl* expression in hemocytes, adult IFMs, and TTCs, confirming that all these three cell types are bona fide sources of Bnl under normal conditions (Figure S6 E-F’). Surprisingly, following injury, *bnl-lacZ* expression was no longer detected in the nuclei of injured muscle fibers, whereas neighboring uninjured fibers retained its expression (Figure S6 G-H’’). Consistent with this observation, muscle-specific depletion of *bnl* using Mef2-Gal4 did not impair tracheal recruitment to damaged fibers (Figure 6 I-L), suggesting that injured muscles are not a major source of Bnl driving this response. In contrast, *bnl-lacZ* reporter expression was readily detected in TTCs recruited to injured muscles (Figure S7 A-B’’). Moreover, knockdown of *bnl* in TTCs impaired tracheal targeting of damaged fibers (Figure S7 C-F), indicating that Bnl produced by the TTCs contributes to tracheal remodelling through an autocrine signalling mechanism. A similar role for autocrine signalling function of Bnl has previously been described during tracheal remodelling in the adult gut [18, 19].

In addition to TTCs, we revealed a marked enrichment of Bnl::GFP^endo^ signal surrounding Pxn+ hemocytes recruited to the damaged fibers (Figure 4 D-E’). This observation was further supported by the detection of *bnl-lacZ* expression in the recruited Pxn+ hemocytes (Figure 4 F-G’’). To assess the functional relevance of hemocyte-derived Bnl to tracheal invasion, we selectively silenced *bnl* in hemocytes using either the *Enh3-Gal4* or *HmlΔ-Gal4* drivers. While hemocyte recruitment to injured muscles was unaffected, *bnl* knockdown significantly reduced tracheal invasion into damaged fibers compared with controls (Figure 4 H-J’’ and K; Figure S7 G-I’’ and J). This signalling between the two cell types aligns with their close anatomical association under homeostatic conditions (Figure 2 A-A’ and [13]). Together, these data indicate that injured muscle fibers themselves do not contribute to the localized accumulation of Bnl following injury. Rather, Bnl is provided by recruited hemocytes and tracheal terminal cells.

Given the critical role of the tracheal system in supporting tissue function [16], we next asked whether hemocyte-mediated tracheal recruitment via Bnl is required to maintain muscle flight performance after injury. To test this, we performed flight assays on uninjured controls, injured flies, and injured flies in which *bnl* was specifically depleted in hemocytes (Figure 4 L-L’). As expected, injured flies displayed reduced flight performance compared with uninjured controls. Importantly, flight performance was further impaired when *bnl* was depleted in hemocytes (Figure 4 L’). These data indicate that hemocyte-derived Bnl contributes to maintaining muscle function following injury, likely through its role in promoting tracheal remodelling (Figure 4 M).

### The receptor FGFR/Btl is essential for injury-induced tracheal guided outgrowth

The finding that Bnl orchestrates tracheal morphogenesis in response to muscle damage suggested in turn that it activates its receptor Btl, on tracheal cells, thereby driving guided outgrowth and branching. To investigate this, we examined Btl expression in adult flight muscles using an endogenously tagged *Btl::cherry^endo^* allele, which faithfully reports Btl expression [31]. Btl::cherry^endo^ was detected along tracheal branches under homeostatic conditions and within tracheal structures invading the injured muscle (Figure 5 A-B’’). To complement this approach, we expressed a membrane-tethered GFP (mCD8-GFP) under the control of a *Btl-Gal4* driver (Figure 5 C-E’). While Btl::cherry^endo^ reflects consistent endogenous expression, *Btl-Gal4* drives a more heterogeneous pattern across tracheal cells (Figure 5 C-C’). Nevertheless, we observed a striking accumulation of tracheal branches labelled by *Btl-Gal4* at injured muscles, reinforcing that Btl-expressing tracheal cells are actively recruited to damaged fibers (Figure 5 D-E’). We then tested whether Btl function is required for proper tracheal pathfinding. To this end, *btl* was specifically silenced in tracheal cells using either *dSRF-Gal4* or *Btl-Gal4* drivers (Figure 5 F-I’’). In both cases, tracheal invasion into injured fibers was significantly impaired compared to the control (Figure 5 J). Notably, the impairment was more pronounced with *dSRF-Gal4*, consistent with its more uniform expression in terminal tracheal cells (Figure 2 F-F’) compared with *Btl-Gal4* (Figure 5 C-C’). Together, these findings show that the Bnl/Btl signalling axis plays a pivotal role in guiding tracheal remodelling after muscle damage. While Btl expression is restricted to tracheal cells, localized Bnl secretion from hemocytes at the damaged fiber refines tracheal pathfinding, ensuring precise and targeted tracheal invasion into injured muscle (Figure 5 K).

**Figure 5.**
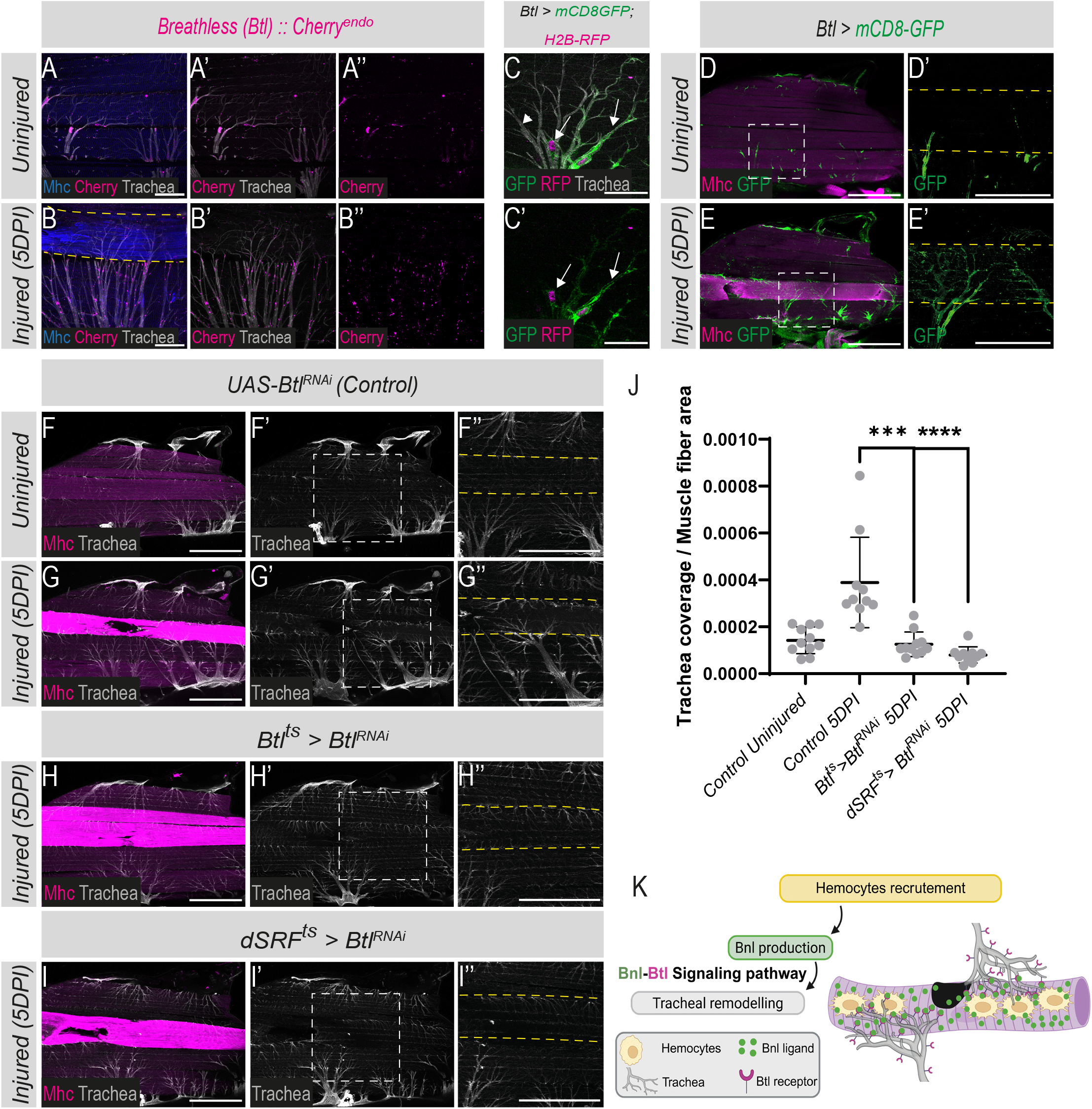
The FGF receptor Breathless (Btl) is expressed along tracheal branches and is required for tracheal recruitment to the damaged muscle. **(A-B’’).** Btl::Cherry^endo^ (magenta) is detected along tracheal branches (gray) associated with indirect flight muscles (IFMs) labelled with Mhc (blue), in either uninjured (A) or injured muscles (B). Scale bars: 50 μm. Dashed yellow outlines indicate injured muscle fibers showing elevated Mhc expression. **(C-C’).** The *Btl-Gal4* driver (green and magenta; *Btl-Gal4, UAS-mCD8GFP; UASH2B-RFP*) is active in a subset of tracheal branches (arrows, C-C’). Arrowheads in C indicate the tracheal branches that do not express the *Btl-Gal4* driver. Scale bars: 25 μm. **(D-E’).** Activity of the *Btl-Gal4* driver (green, *Btl-Gal4, UAS-mCD8GFP*) in uninjured condition (D-D’) and at 5DPI (E-E’). Muscles are labelled with Mhc (magenta). Scale bars: 200 μm. **(F-I’’).** Downregulating *btl* in the tracheal cells with either *Btl-Gal4* driver (*Btl-Gal4; UAS-Btl^RNAi^*, H-H’’) or *dSRF-Gal4* driver (*dSRF-Gal4, UAS-Btl^RNAi^*, I-I’’) prevents tracheal invasion to the damaged fiber. **(J).** Quantification of trachea coverage (****p <0.0001, **p=0.0053, Student’s t-test; n > 10 heminota per condition). Scale bars: 200 μm. **(K).** Schematic model of hemocyte-driven Bnl activates tracheal cells recruitment via Bnl/Btl signalling pathway.

### ECM provided by the hemocytes retains Bnl ligand at injured muscles

Although Bnl is a diffusible molecule, it is specifically retained at the injured fiber (Figure 3 K’-K’’), raising the question of what restricts its spatial distribution. Previous study has shown that Collagen IV, a principal structural component of the muscle ECM, is critical for retaining diffusible signals within defined tissue regions [32, 33]. We therefore hypothesized that ECM components may similarly act to confine Bnl to the damaged fiber. Hemocytes are known to be major producers of ECM components during both *Drosophila* development and post-developmental stages [34, 35], raising the possibility that they could supply Collagen IV in response to adult muscle injury. To test this, we analysed Collagen IV distribution in adult flight muscles using the Vkg::GFP line (Figure 6 A-B’’). Under homeostatic conditions, we observed a basal level of Vkg::GFP surrounding muscle fibers and along tracheal branches (Figure 6 A’), and we found that hemocytes express Collagen IV (Figure 6 A’’). Five days after injury, Vkg::GFP was markedly enriched in damaged fibers compared with intact muscles and uninjured controls (Figure 6 A-B and A’’-B’’). To assess the relationship between hemocytes recruitment and ECM remodelling, we examined Collagen IV distribution across multiple time points post-injury (1-20 DPI) and compared this to hemocyte recruitment dynamics (Figure 6 C-D’’ and Figure S8 A-F). Collagen IV accumulation closely correlated with the presence and distribution of hemocytes. Notably, regions where hemocytes were evenly distributed along the injured fiber displayed stronger and more uniform Vkg::GFP signal. Consistently, downregulation of Collagen IV in the hemocytes significantly reduce Vkg::GFP signal at the injured muscles (Figure S8 G-I). Together, these data indicate that hemocytes make a predominant contribution to the local Collagen IV remodelling response.

**Figure 6.**
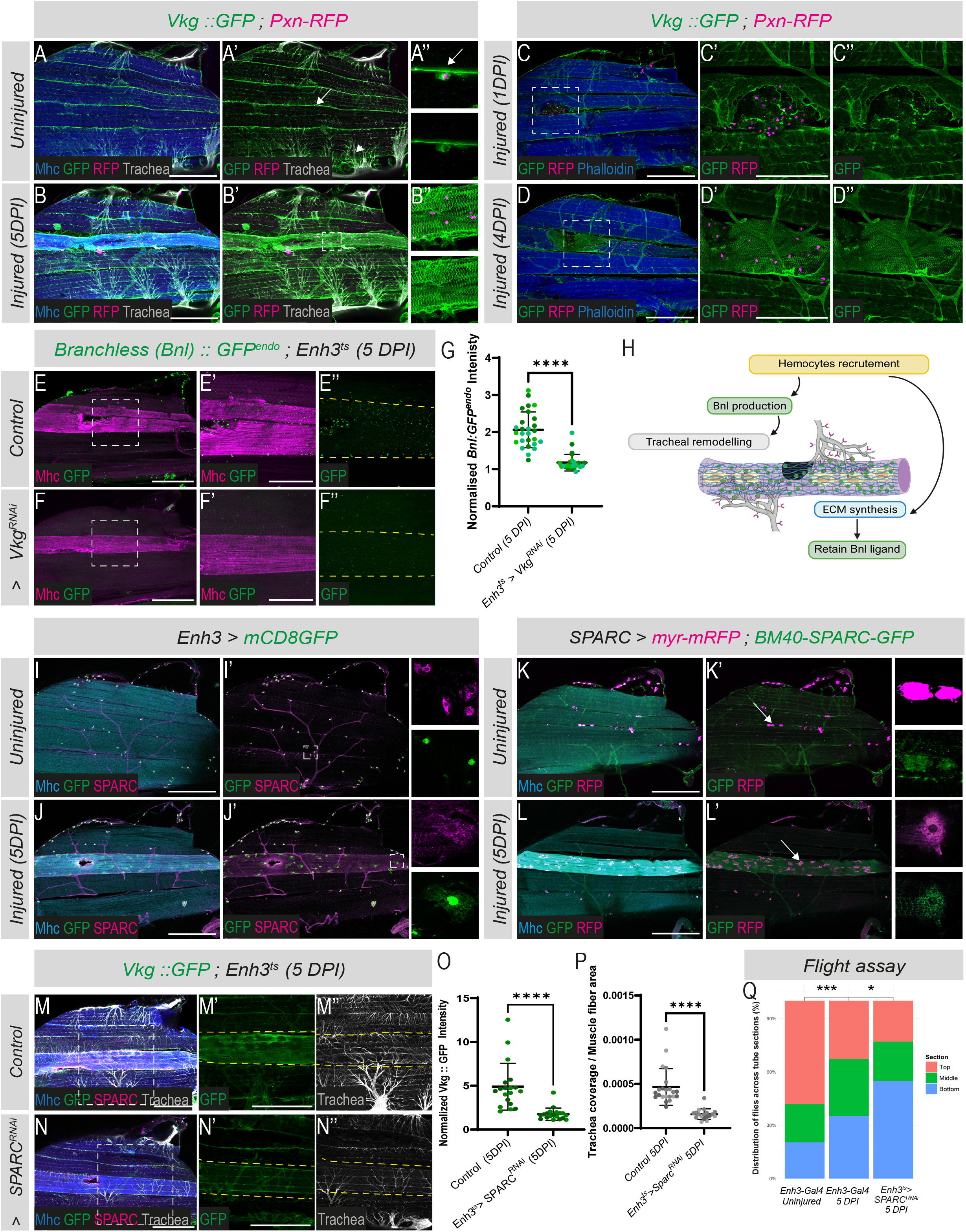
Hemocytes deposit ECM components at injury muscles to constrain Bnl to damaged muscle fibers and maintain muscle function. **(A-A’’).** Collagen IV (green, *Vkg::GFP*) is detected at basal levels around muscle fibers (arrow, A’), tracheal branches (arrowheads, A’), and hemocytes (magenta, *Pxn-RFP*; arrow, A’’) in uninjured conditions. **(B-B’’).** At 5DPI, Collagen IV is strongly enriched on the injured muscle fibers, labelled with Mhc (blue). **(C-D’’).** Hemocytes recruitment (magenta, *Pxn-RFP*) coincides with Collagen IV (green, *Vkg::GFP*) distribution on the injured muscle, at 1DPI and 4DPI. Muscles are labeled with phalloidin (blue). Scale bars: 200 μm. **(E-F’’).** The endogenous Bnl (green, *Bnl::GFP^endo^*) is enriched along damaged muscle fibers (magenta, Mhc) and is markedly reduced upon hemocyte-specific depletion of Collagen IV (*Enh3-Gal4, UAS-Vkg^RNAi^*). **(G).** Quantification of normalized Bnl::GFP^endo^ intensity in injured fibers (****p<0.0001, Student t-test, n >27 heminota per condition from three independent replicates). **(H).** Schematic model of hemocyte-mediated ECM delivery required to confine Bnl to injured muscle fibers. **(I-I’).** SPARC (magenta) is detected in the IFMs associated hemocytes visualized by *Enh3-Gal4* driver (green, *Enh3-Gal4, UAS-mCD8GFP*). **(J-J’).** SPARC expression is increased in injured fibers compared to uninjured condition (I-I’). **(K-K’).** Adult hemocytes (magenta; *SPARC-Gal4, UAS-myr-RFP*) express BM40-SPARC-GFP (green). **(L-L’).** BM40-SPARC-GFP produced by adult hemocytes is broadly distributed along the injured muscle fibers. **(M-N’).** Downregulation of *SPARC* in adult hemocytes (*Enh3-Gal4 > UAS-SPARC^RNAi^*) reduces Collagen IV enrichment (green, *Vkg::GFP*) and impairs tracheal recruitment in injured fiber (gray, autofluorescence), as quantified in panels **(O)** and **(P)** respectively (O. ****p<0.0001, Student t-test, n = 19 heminota per condition from two independent replicates, P. ***p<0.001, Student t-test, n = 18 heminota per condition from two independent replicates). **(Q).** Quantification of the proportion of flies landing in the three flight-assay zones (Top, Middle, and Bottom) under the following conditions: Control Uninjured, Control Injured (5 DPI), and *Enh3-Gal4 > UAS-SPARC^RNAi^*Injured (5 DPI) (***p < 0.001, *p < 0.05, Chi-square test; n ≥ 100 flies per condition from three independent biological replicates).

To determine whether Collagen IV accumulation reflects a general remodelling of the ECM or the selective enrichment of specific components, we next analysed the distribution of Laminin A. Laminin A is another major constituent of the muscle basement membrane that has been reported to associate with adult IFMs and their tracheal cells [24]. In contrast to Collagen IV, Laminin A was not enriched at injured fibers following damage (Figure S8 J-K). Furthermore, muscle-associated hemocytes did not express Laminin A in either uninjured or injured adult muscles (Figure S8 J-K). These findings indicate that injury-induced ECM remodelling is selective and involves the preferential enrichment of specific basement membrane components, including Collagen IV.

To test whether hemocyte-supplied Collagen IV is necessary for Bnl retention at injured fibers, we knocked down Collagen IV specifically in hemocytes and assessed Bnl::GFP^endo^ localization (Figure 6 E-F’’ and Figure S9 A-C). In control flies, Bnl signal is enriched along the injured fiber; however, this enrichment was significantly reduced following hemocyte-specific Collagen IV depletion (Figure 6 E-F’’ and G). Importantly, hemocytes recruitment to damaged fibers was not affected by Collagen IV depletion (Figure S9 A-D), indicating that Collagen IV is not required for immune cell recruitment to injured fibers. These data show that hemocyte-derived Collagen IV is required to spatially confine Bnl to the damaged fibers (Figure 6 H).

We next asked whether loss of Bnl confinement following hemocyte-specific Collagen IV depletion affects tracheal remodelling after muscle injury. Absolute tracheal enrichment at injured fibers was comparable between control and Collagen IV–depleted conditions (Figure S9 E). However, the ratio of tracheal coverage at injured muscles relative to neighbouring intact muscles was significantly reduced when Collagen IV was depleted in hemocytes (Figure S9 E’). These data indicate that loss of ECM induces Bnl signal to spread beyond the injury site, thereby provoking non-specific tracheal remodelling in adjacent, intact muscles.

### SPARC promotes Collagen IV deposition and tracheal remodelling following muscle injury

Incorporation of Collagen IV into the extracellular matrix (ECM) requires accessory molecules for proper deposition and stabilization. SPARC is a collagen-binding glycoprotein essential for Collagen IV diffusion, stabilization, and incorporation into basement membranes [36, 37]. We therefore investigated whether hemocytes also supply SPARC in response to muscle damage. We first examined SPARC localization in hemocytes associated with adult IFMs. Hemocytes, identified using *Enh3-Gal4* driver, were positive for SPARC (Figure 6 I-I’). Importantly, following muscle injury, SPARC levels were elevated along the injured fiber compared with controls, mirroring the distribution of Collagen IV (Figure 6 I-J’). These observations suggest that SPARC is locally produced by hemocytes in adult muscle. To further confirm the cellular origin of SPARC enrichment in injured muscles, we employed the SPARC-Gal4 driver, a Gal4 trap line inserted into the endogenous SPARC locus, to drive membrane-targeted RFP (myr-RFP) expression in flies also carrying a BM40-SPARC-GFP fosmid reporter (Figure 6 K-L’ and Figure S9 F-G’). This dual-reporter approach allowed simultaneous visualization of SPARC (via BM40-SPARC-GFP) and identification of SPARC-producing cells (via SPARC-Gal4 > myr-RFP). In uninjured muscle, BM40-SPARC-GFP and SPARC-Gal4-driven myr-RFP were co-expressed in hemocytes. Following injury, BM40-SPARC-GFP was broadly enriched along the damaged fiber, resembling endogenous SPARC protein, whereas SPARC-Gal4 activity remained restricted to hemocytes and was not detected in muscle nuclei (Figure 6 L-L’ and Figure S9 G-G’). Together, these data demonstrate that recruited hemocytes are the primary source of SPARC at injured muscle fibers. We then tested whether hemocyte-derived SPARC is required for Collagen IV enrichment and tracheal remodelling. While silencing *sparc* in hemocytes does not prevent their recruitment to injured muscles (Figure S9 J-J’’’), it is sufficient to prevent both Collagen IV accumulation and tracheal remodelling at the injured fiber (Figure 6 M-N’’ and O-P). Finally, to evaluate the functional importance of SPARC in maintaining muscle function, we performed flight assays and found that injured flies lacking hemocyte-derived SPARC exhibited significantly reduced flight ability compared with controls (Figure 6 Q). These results support a functional role for hemocyte SPARC in preserving muscle performance following damage. We also note that we cannot completely exclude an additional effect linked to altered immune function. Altogether, these results show that hemocyte-derived SPARC is essential for ECM assembly, tracheal remodelling, and functional recovery following muscle injury. They also highlight that hemocytes act as multifunctional signalling hubs, simultaneously delivering trophic, structural, and guidance cues required to coordinate efficient muscle repair.

## Discussion

The maintenance of tissue function following damage requires the coordinated integration of immune, vascular, and extracellular matrix responses, yet how these processes are mechanistically linked remains poorly understood. Here, we identify a signalling program in which hemocytes connect muscle damage to tracheal remodelling and extracellular matrix organization. We show that hemocytes respond to injury by locally providing the FGF ligand Branchless to guide tracheal outgrowth toward damaged muscle, while simultaneously supplying extracellular matrix components that promote Bnl retention. Together, these processes contribute to the stabilization of muscle function following injury and likely limit further functional decline. Our findings therefore identify hemocytes as central coordinators of the response to damage in adult *Drosophila* muscle (Figure 7).

**Figure 7.**
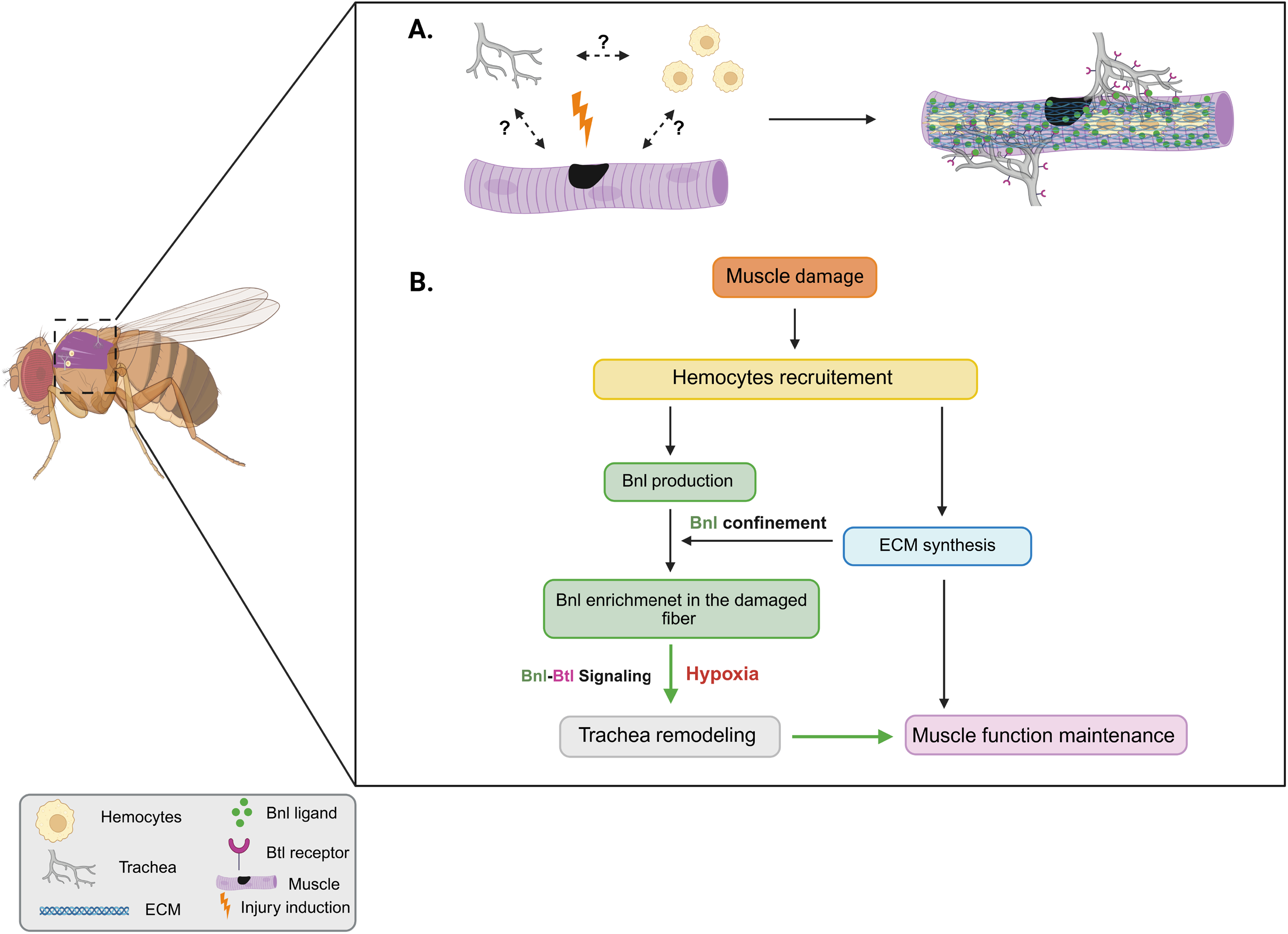
Hemocyte-muscle-tracheal intercellular signaling preserve muscle function following acute damage. **(A).** Schematic representation of the hemocyte-muscle-tracheal signaling module activated in response to muscle injury. **(B).** Upon muscle damage (yellow lightning symbol), hemocytes are rapidly recruited to the injured fiber, where they secrete the FGF-like ligand Branchless (Bnl), which activates its receptor Breathless (Btl) on tracheal cells. Bnl-Btl signaling drives tracheal remodeling and directional outgrowth toward the injured muscle. In parallel, hemocytes deposit extracellular matrix (ECM) components that retain Bnl and stabilize trachea-muscle attachments. These coordinated cellular mechanisms preserve flight performance.

### Immune-derived FGF acts as an intercellular signalling cue to coordinate vascular remodelling in response to muscle damage

FGF signalling contributes to multiple aspects of vertebrate myogenesis and muscle regeneration [38–41]. For example, FGF2 stimulates myoblasts proliferation while suppressing differentiation *in vitro*, whereas FGF6 is required to maintain full regenerative capacity and MuSC pool size [42, 43]. However, whether FGF signalling also mediates vascular remodelling during vertebrate muscle repair, as it does in other regenerative contexts, remains unclear. One explanation for this gap may be the extensive functional redundancy of the vertebrate FGF family, which comprises multiple ligands with overlapping expression patterns and signalling activities. This redundancy complicates precise genetic dissection of individual FGF functions during regeneration. Moreover, the cellular sources of FGF ligands during muscle regeneration remain poorly defined [43], further limiting targeted genetic ablation strategies.

Here, we show that following muscle damage, hemocytes are rapidly recruited to and spread over the injured muscle surface, where they secrete the FGF ligand Branchless (Bnl) to promote localized tracheal outgrowth and precise branch targeting (Figure 7). This mode of immune-driven Bnl delivery contrasts with the prevailing model in which the tissue requiring vascular or tracheal infiltration is itself thought to produce the FGF signal [29, 44]. The identification of a non–cell-autonomous source of Bnl during the muscle injury response is therefore unexpected and raises an important question: why does the injured muscle fiber not itself express Bnl to attract tracheal branches? One possible explanation is the need to tightly restrict tracheal growth. Bnl is a potent chemoattractant, and its widespread expression by muscle fibers could lead to excessive or ectopic tracheogenesis, resulting in inappropriate branch recruitment and disruption of normal tracheal network architecture. In contrast, hemocyte enrichment at damaged muscles creates a localized source of Bnl, resulting in ligand accumulation that is not observed at other physiological sites occupied by hemocytes. Such spatially restricted delivery would ensure that tracheal remodelling occurs specifically at sites of injury, where oxygen delivery is most acutely required. Accordingly, basal Bnl expression in healthy muscle fibers may support existing tracheal associations, whereas local Bnl enrichment following injury promotes additional tracheal recruitment. Furthermore, the sequential recruitment of hemocytes, initially to the injury site and later along the entire damaged fiber, suggests a mechanism by which Bnl signalling can be progressively deployed to support tracheal invasion of the injured muscle.

The nature of the signals governing hemocyte recruitment to injured muscles remains unknown. Previous studies in *Drosophila* have shown that tissue damage induces the production of reactive oxygen species (ROS), which can promote hemocytes activation and migration toward wounded tissues [45]. Additional damage-associated cues, including DAMPs [46] and inflammatory signaling molecules [47] may also contribute to this process. Although we detected activation of the hypoxia reporter in a subset of recruited hemocytes (Figure S5 A-B’’’), disruption of hypoxia signaling did not affect hemocyte accumulation at injured muscles (Figure S5 H-J’’’ and L), suggesting that hypoxia is not a major driver of the initial recruitment process. Future studies will be required to identify the signals that attract hemocytes to damaged muscle fibers and to determine how these cues are integrated with local hypoxic and ECM remodelling responses.

### Local ECM delivery directs vascular remodelling toward damaged muscle

The ECM is essential for muscle integrity and function [48]. However, how ECM remodelling is coordinated with other cellular processes in response to muscle damage remains incompletely understood. In vertebrates, fibro-adipogenic progenitors (FAPs) are the principal source of ECM within muscle tissue [49]. *Drosophila* lack a dedicated FAP population. During development, hemocytes are one of the principal sources of ECM components in flies [34, 50, 51]. Here, we show that adult hemocytes also produce key ECM components, including Collagen IV and SPARC, that are required for proper response to muscle damage. Nevertheless, the requirement for hemocyte SPARC and Collagen IV in response to damage identifies hemocytes as a major functional source of ECM during this process. These findings suggest that Drosophila hemocytes share functional similarities with mammalian FAPs and may partially compensate for their absence, pointing to an evolutionarily flexible strategy for muscle maintenance and repair in organisms with fewer specialized cell types. We note, however, that we do not exclude a potential contribution of the fat body to ECM delivery following muscle injury.

We find that hemocyte-derived ECM forms a specialized local microenvironment in which FGF distribution is spatially constrained. Interestingly, a conceptually similar mechanism has been described during *Drosophila* gonad development, where “companion” hemocytes locally deposit ECM components to support tissue morphogenesis [33]. In this system, the ECM has been shown to physically interact with Dpp and restrict its diffusion [32]. Whether the ECM physically interacts with Bnl or instead acts primarily as a diffusion barrier in response to muscle damage remains to be determined. Overall, these findings provide a mechanistic example of how extracellular matrix remodelling can be coordinated with vascular remodelling to support tissue maintenance following muscle injury.

### Do Drosophila repair or regenerate their flight muscles?

The IFMs generate the high-frequency wing beats that enable flight, a behavior essential for predator avoidance, foraging, and mating. Given their vital function, IFMs must be maintained in optimal condition to ensure sustained performance throughout the animal’s lifespan. This need for resilience is also underscored by the observation that over 30% of wild-caught flies exhibit physical injuries, frequently located in the thorax and abdomen [52]. Although a rare population of satellite-like cells has been identified in adult flies, the regenerative capacity of adult *Drosophila* muscles remains debated [53]. A major limitation has been the lack of appropriate molecular markers to monitor both muscle regeneration and muscle stem cell (MuSC) activity. In our study, we identified a specific Myosin Heavy Chain (Mhc) isoform that is consistently detected at elevated levels in damaged fibers. However, unlike in vertebrates, where MyHC-emb expression returns to baseline following regeneration, Mhc detection in *Drosophila* remains elevated even 20 days post-injury (Figure S1 B-D). This makes Mhc a sensitive marker for identifying injury, but less useful for assessing complete muscle recovery. Adult MuSCs have been identified via the transcription factor Zfh1/ZEB and its reporter Enh3 [9, 10]. Zfh1 is required for MuSCs maintenance, an equivalent role of ZEB was subsequently shown in mice [54]. Interestingly, both Zfh1 and ZEB are expressed in the muscle associated immune cells [9, 54]. In our study, we found that following muscle injury, 100% of the cells recruited to damaged fibers are both Enh3⁺ and Pxn⁺ (a plasmatocytes marker), while being negative for the muscle-specific marker Mef2 (Figure S3 E-F’). This strongly suggests that hemocytes are the predominant, if not exclusive cell population migrating to injured muscle. These findings suggest that adult *Drosophila* flight muscles possess limited intrinsic regenerative capacity and may instead rely on compensatory mechanisms such as increased oxygenation through tracheal remodelling and structural reinforcement via extracellular matrix (ECM) deposition-to cope with muscle damage. One possible advantage of such a strategy is that it may help stabilize injured fibers and preserve the integrity of the flight muscle apparatus as a whole. This could limit collateral damage to neighboring, unaffected muscle fibers. Such a response is more consistent with tissue repair than with true regeneration. Regeneration implies the restoration of the original tissue architecture through replacement of damaged cells and reconstruction of the tissue. In contrast, repair encompasses a broader range of adaptive responses that preserve tissue integrity and function following damage, without necessarily restoring the original structure [55]. This is consistent with the broader idea that short-lived species prioritize immediate survival over long-term regenerative investment [56]. In such organisms, evolution favors fast and efficient repair mechanisms that limit functional decline and preserve organismal fitness over complex, energy-intensive regeneration programs.

## Materials and Methods

### Drosophila melanogater strains and genetics

All *Drosophila melanogaster* stocks were grown on standard medium at 25°C. The following stains were used: *w^118^*as wild type (wt), *Pxn-RFP* [57], *HmlΔ-Gal4* (BL#30140)*, Enh3-Gal4* [9]*, Mhc-Iso-K, L, M (fTRG500)* and *Mhc-Iso-A, F, G (fTRG519)* [24], *weeP26* [23], *Bnl::GFP^endo^ and Btl::cherry^endo^* [31], *bnl-LacZ* (BL#11704), *dSRF-GAL4* (BL#25753), *Btl-Gal4* (BL#78328), *UAS-Btl^RNAi^*(BL#435544), *UAS-Bnl^RNAi^* (BL#34572), *UAS-Bnl::GFP* [58], *Vkg::GFP* (BL#98343), *BM40-SPARC-GFP* (Sarov et al., 2016), *SME-Gal4* (VT 0576871), *UAS-H2B-RFP* [27], *UAS-mCD8GFP* (BL#5137), *UAS-myr-mRFP* (BL#7118), SyHREns [30], *UAS-Sima^RNAi^* (BL#26207), *SPARC-Gal4* (BL#77473), *UAS-SPARC^RNAi^* (BL#40885), *UAS-Vkg^RNAi^* (BL#50895), *Laminin A::GFP* (VDRC #318155). *Naa30A* deletion males were obtained from crosses between *Naa30A^Δ74^/FM0* females and either UAS-mCD8GFP; Enh3-Gal4 (Figure 1) or Mhc-Iso-K, L, M (Figure S1) males. RNAi experiments were carried out at 29°C. To achieve temporal control of gene expression in adult tissues, the tubGal80^ts^ was employed, allowing restriction of Gal4 activity to a specific time window. Fly crosses were initially maintained at 18°C, and after adult eclosion, the progenies were transferred to 29°C, where they remained until dissection. An exception was made for the Bnl knockdown experiments shown in Figure 4 J-J’’, which were performed without the tubGal80^ts^ transgene.

### Generation of SME-FRT-stop-FRT-Gal4 fly line and flip-out induction procedure

SME (Shavenbaby Muscle Enhancer) is one of several enhancers of the *shavenbaby* gene (VT057071-Gal4). It is active in larval muscle progenitors (AMPs) (Figure S6 A-A’) and remains active in a subpopulation of localized muscle nuclei in the adult indirect flight muscles (IFMs) (Figure S6 B-B’). To generate the SME-FRT-stop-FRT-Gal4 construct, the cis-regulatory element from the original *shavenbaby* enhancer construct was used to replace the *pAct* promoter in the pAct-FRT-stop-FRT3-FRT-FRT3-Gal4 vector (Addgene plasmid #52889). The resulting construct was integrated into the attP site at cytological location 51D on chromosome II via embryo injection into vas-int; attP-51D flies. To induce localized overexpression of Bnl::GFP in adult IFM nuclei, hsflp; FRT-SME-stop-FRT-Gal4/CyO; tubGal80^ts^ females were crossed to UAS-Bnl::GFP males. Progeny were raised at 18°C and subjected to a 2-hour heat shock at 37°C in young adulthood (Figure 4 A-C’).

### Immunohistochemistry and in situ hybridization

Preparation and immunofluorescence of wing discs and adult muscles were performed as previously described in [59]. The following primary antibody were used: Chicken anti-βGal (1:100, Abcam, ab9361), Chicken anti-GFP (1:300, Abcam, ab13970), Mouse anti-Mhc (1:200, DSHB, 3E8-3D3-s), Mouse anti-dSRF (1:20, a gift from Mark A. Krasnow, Stanford university California, USA), Rabbit anti-Mef2 (1:500, DSHB, Mef2-p), Rabbit anti-PH3 (1:100, Cell Signaling Technology, #9713), Rabbit anti-SPARC (1:500, a gift from Maurice Ringuette, University of Toronto, Canada), Rabbit anti-Zfh1 (1:5000, a gift from Ruth Lehmann, New York, USA), Rat anti-RFP (1:800, Chromotek, 5F8-20), Rabbit anti-cleaved *Drosophila* Dcp-1(1:400, Cell Signaling Technology, #9578), Alexa-conjugated Phalloidin (1:200, Thermo fisher scientific life technologies). *mef2* single-molecule fluorescence in situ hybridization (smFISH) experiments were performed as described in [59].

### Induction of flight muscle injury

Flight muscle injury was induced following previously described protocol [60]. Adult flies (2-3 days post-eclosion) were anesthetized and positioned laterally under a stereomicroscope. A single, localized stab wound was inflicted manually using a fine stainless-steel needle (Minutien Pins, 0.1 mm diameter; #26002–10) to minimize tissue damage and avoid extensive injury. Age-matched, uninjured flies served as controls.

### Flight assay

Flight ability was assessed using a 1-m-long cylindrical flight column internally lined with paper coated with a thin layer of glue. The adhesive sheet was divided evenly into three longitudinal sections corresponding to the Top, Middle, and Bottom regions of the tube (Figure S1). For each experimental condition, an equal number of flies (approximately 20–30) was released into the tube. After all flies had landed, the paper was removed, flattened, and imaged for analysis.

Image quantification was performed semi-automatically using a custom R script, Fly-Spy (available at https://github.com/Olivier-Josse/Fly-spy). Images were first converted to grayscale and digitally divided into the three predefined regions. High-intensity spots corresponding to individual flies were identified by applying a manually adjusted threshold to ensure accurate segmentation. The number of flies detected in each region was quantified and expressed as a percentage of the total recovered flies. The distribution of flies across the three regions—reflecting high, intermediate, or low/absent flight ability—was compared between experimental conditions using a chi-square (χ²) test. Results are presented as stacked bar histograms

### Microscopy and data analysis

Samples were imaged using a Leica SP8 microscope (BIOSIT, University of Rennes) at 20×, 40×, or 63× magnification and a resolution of 1024×1024 pixels. Images were processed with ImageJ and assembled using Adobe Illustrator. To quantify fluorescence intensity, all samples were processed and analyzed in parallel under identical conditions. Confocal images were acquired using the same laser settings across all samples. Sum slice projections were generated from each confocal stack. Regions of interest (ROIs) were manually defined based on protein expression, and mean fluorescence intensities were measured within each ROI. These values were then normalized to background levels measured in comparable areas of the same sample.

### Quantification of tracheal coverage and enrichment

Tracheal branches were visualized either by their intrinsic autofluorescence using a 405 nm UV laser [15] or by the expression of membrane-bound GFP driven by a tracheal-specific Gal4 driver (UAS-mCD8GFP). Tracheal coverage was quantified from confocal stacks acquired at 20X magnification using the approach described by Perochon et al. (2021) [18], with minor modifications. To ensure valid comparisons between conditions, equivalent z-stack depths were analysed in all samples. Fibers #1 and #5 (Figure 1 G’ and H’) were excluded from the analysis because their proximity to the primary tracheal trunks originating from the dorsal and ventral air sacs. Maximum-intensity projections were generated from each confocal stack, tracheal branches were segmented by thresholding, and the resulting images were skeletonized using the ImageJ “Skeletonize” plugin. Skeleton pixel intensity was then measured within injured muscle fibers and their corresponding contralateral control fibers. To account for potential differences in tissue size between conditions, all tracheal measurements were normalized to the corresponding muscle area. For Figure 2E, square regions of interest (ROIs) were placed at the injury site, as well as in proximal and distal regions, to assess local tracheal enrichment. Hemocyte and dSRF-positive cell numbers were quantified manually using ImageJ.

Graphs and statistical analyses were performed using GraphPad Prism. Pair comparisons were statistically tested with a two-sided Student’s t-test. NS, not significant (p>0.05); *p < 0.05, **p < 0.01, ***p < 0.001 ****p< 0.0001. Figures 4L, 4M,, 5K, 6H, 7, S1A and S6C were created using BioRender.com (biorender.com).

## Acknowledgments

We would like to thank Jennifer Zanet, Cedric Polesello, Sougata Roy, Ruth Lehmann, Maurice Ringuette, Jean-Paul Vincent, and Mark A. Krasnow, as well as the Bloomington Drosophila Stock Center, the Vienna Drosophila Resource Center (VDRC), and the Developmental Studies Hybridoma Bank for providing *Drosophila* strains and antibodies. We are grateful to Juila Cordero, Lucas Waltzer, Laetitia Bataille, Alain Vincent, and Michèle Crozatier for their critical reading of the manuscript, and to other members of H.B’s laboratory for valuable discussions. We also thank Emmanuel Gallaud for helpful input during the early phases of the project. We acknowledge Xavier Pinson from the Microscopy Rennes Imaging Center (MRic, BIOSIT, Biogenouest) for technical assistance, and Cyrille Surbled for fly stock maintenance and technical help. Work in the Boukhatmi lab was supported by an AFM-Téléthon Trampoline Grant (#23108), ATIP-Avenir (CNRS), and ANR GenepiMuSC (ANR-23-CE13-0006). N.A. was supported by an AFM-Téléthon PhD fellowship (#23846) and an FRM fellowship (FDT202404018608). E.L was supported by an AFM-Téléthon PhD fellowship (#29992). Work in the Cordero lab was supported by a Sir Henry Dale Fellowship (Wellcome Trust and Royal Society, 104103/Z/14/Z), a Wellcome ISSF Returner Scheme (204820/Z/16/Z), and a Tenovus Fellowship (Tenovus/S22-28) to J.P. The funders had no role in study design, data collection and analysis, decision to publish, or preparation of the manuscript.

## Supplementary figures legends

**Figure S1.**
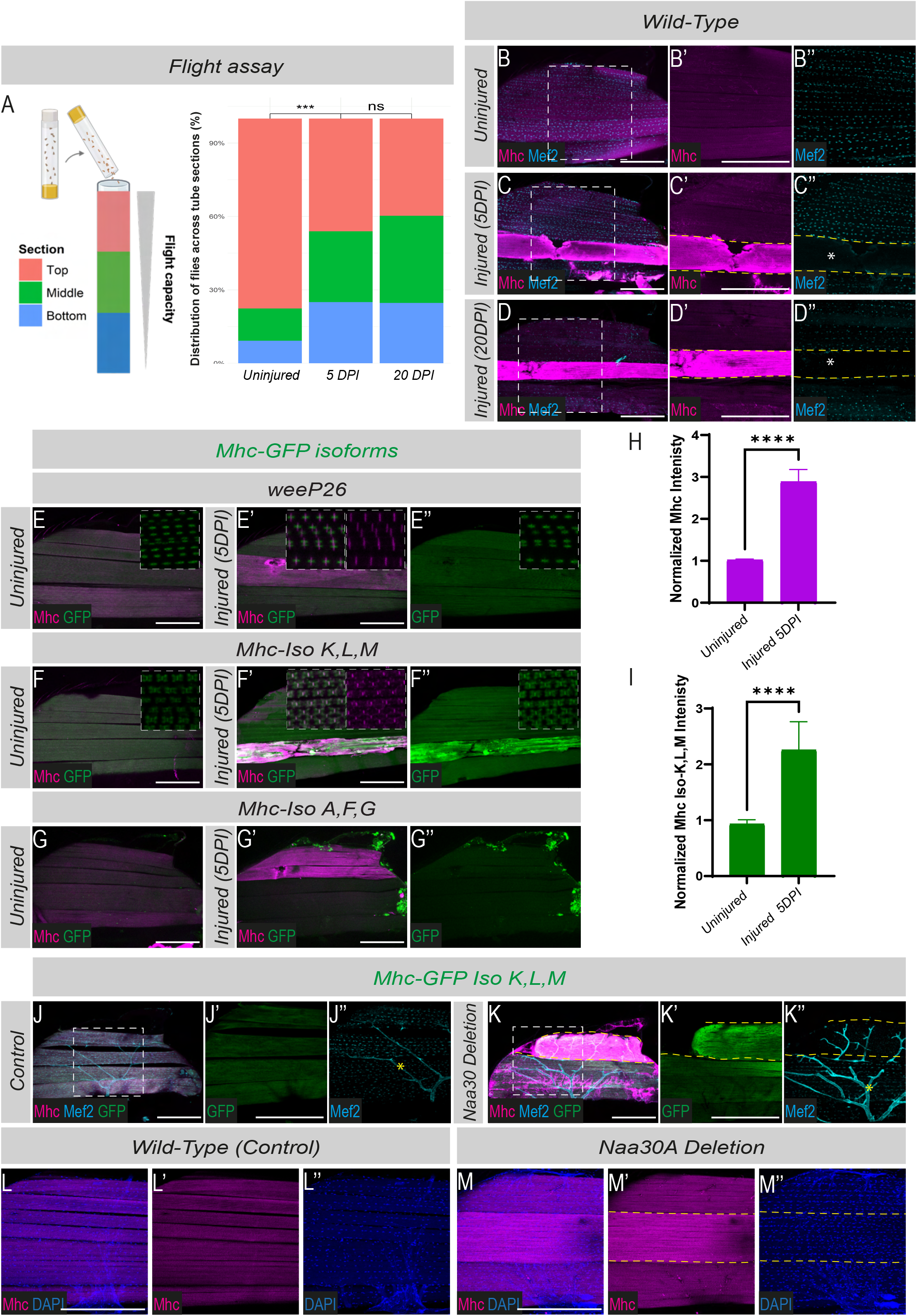
Enhanced Mhc detection marks damaged muscle fiber. **(A)** (Right) Schematic representation of the flight assay used to evaluate flight performance. (Left) Quantification of fly distribution across the three flight-assay zones under uninjured conditions and at 5 and 20 days post-injury (DPI). Bars represent the percentage of flies landing in each zone (***p < 0.001, Chi-square test; n ≥ 80 flies per condition from two independent biological replicates). **(B-D’’).** IFMs stained for Mhc (magenta) and Mef2 (cyan) show enhanced Mhc staining and reduced nuclear Mef2 signal in the injured fiber (asterisks) at 5 (C’-C’’) and 20 DPI (D’-D’’) compared to uninjured IFMs (B’-B’’). (B’, C’ and D’). 2X magnification of the boxed regions in B, C and D. Scale bars: 200 μm. **(E-E’’).** Expression of the embryonic Mhc isoform (*weeP26*, green) in uninjured (E) and injured (E’-E’’) IFMs. Injury does not alter weeP26 detection (E’-E’’). **(F-F’’).** The shorter Mhc isoforms (*Mhc-Iso-K, L, M (fTRG500)*; green) colocalize with anti-Mhc staining (magenta) and are highly detected in injured muscle (F’-F’’). **(G-G’’).** The longer Mhc-GFP isoforms (*Mhc-Iso-A, F, G (fTRG519)*; green) are not expressed in adult IFMs and are not enhanced in response to muscle injury (E’-E’’). Scale bars: 200 μm. (H). Quantification of normalized Mhc intensity. **(I).** Quantification of normalized Mhc Iso-K,L,M intensity. (****p<0.0001, Student t-test, n = 12 heminota per condition from two independent replicates). **(J–K’’).** In Naa30A mutant IFMs, detached muscle fibers show increased detection of both endogenous Mhc (magenta, K) and the short Mhc-GFP isoforms (green, K’), accompanied by a loss of Mef2 expression (cyan, K’’). Scale bars: 200 μm. Yellow asterisks in panels J’’ and K’’ indicate non-specific signal detected in motor neurons. **(L-M’’).** Muscle nuclei visualized with DAPI (blue) in control muscles (L-L’’) and *Naa30A* mutants (M-M’’). Dashed yellow outlines indicate damaged muscle fiber showing enhanced Mhc signal (magenta). Scale bars: 200 μm

**Figure S2.**
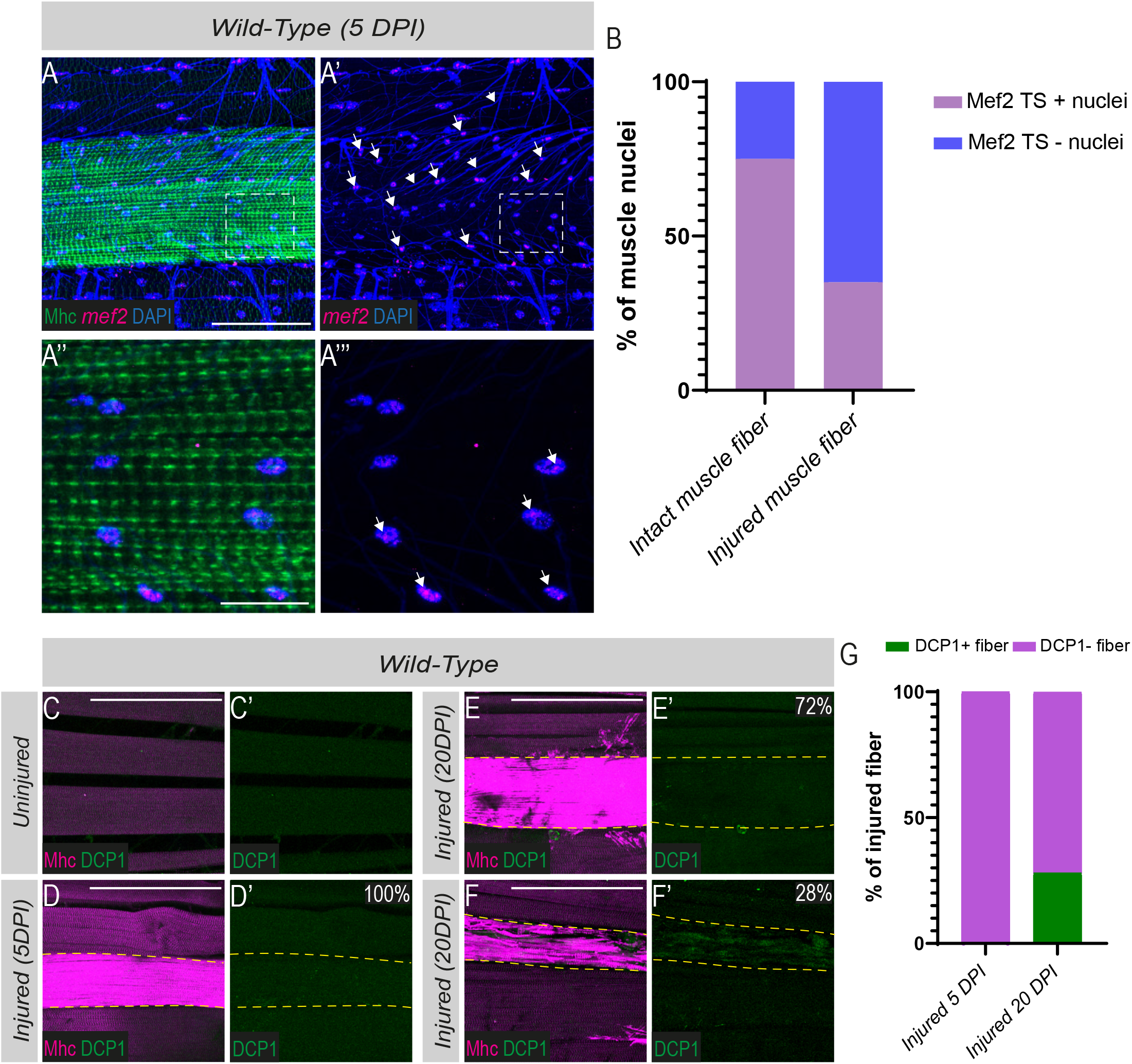
**(A–A’’’).** Active *mef2* transcription sites (magenta, arrows) detected by smFISH in nuclei of injured muscle fibers. Muscles are labelled with Mhc (green), and nuclei are stained with DAPI (blue). Elevated Mhc detection identifies the injured muscle fiber. Arrowheads indicate the tracheal network (autofluorescence). **(B).** Quantification of the percentage of muscle nuclei containing active *mef2* transcription sites (mef2 TS⁺ nuclei) in injured and intact muscle fibers. A total of 435 nuclei from intact muscles and 353 nuclei from injured muscles were analysed. **(C-F’).** Adult IFMs stained for Mhc (magenta) and Dcp1 (green) under uninjured conditions (C-C’), at 5 DPI (D-D’) and 20 DPI (E-F’). **(G).** Quantification of Dcp1-positive injured muscle fibers proportion (n = 30 heminota from two independent replicates). Scale bars: 100 μm.

**Figure S3.**
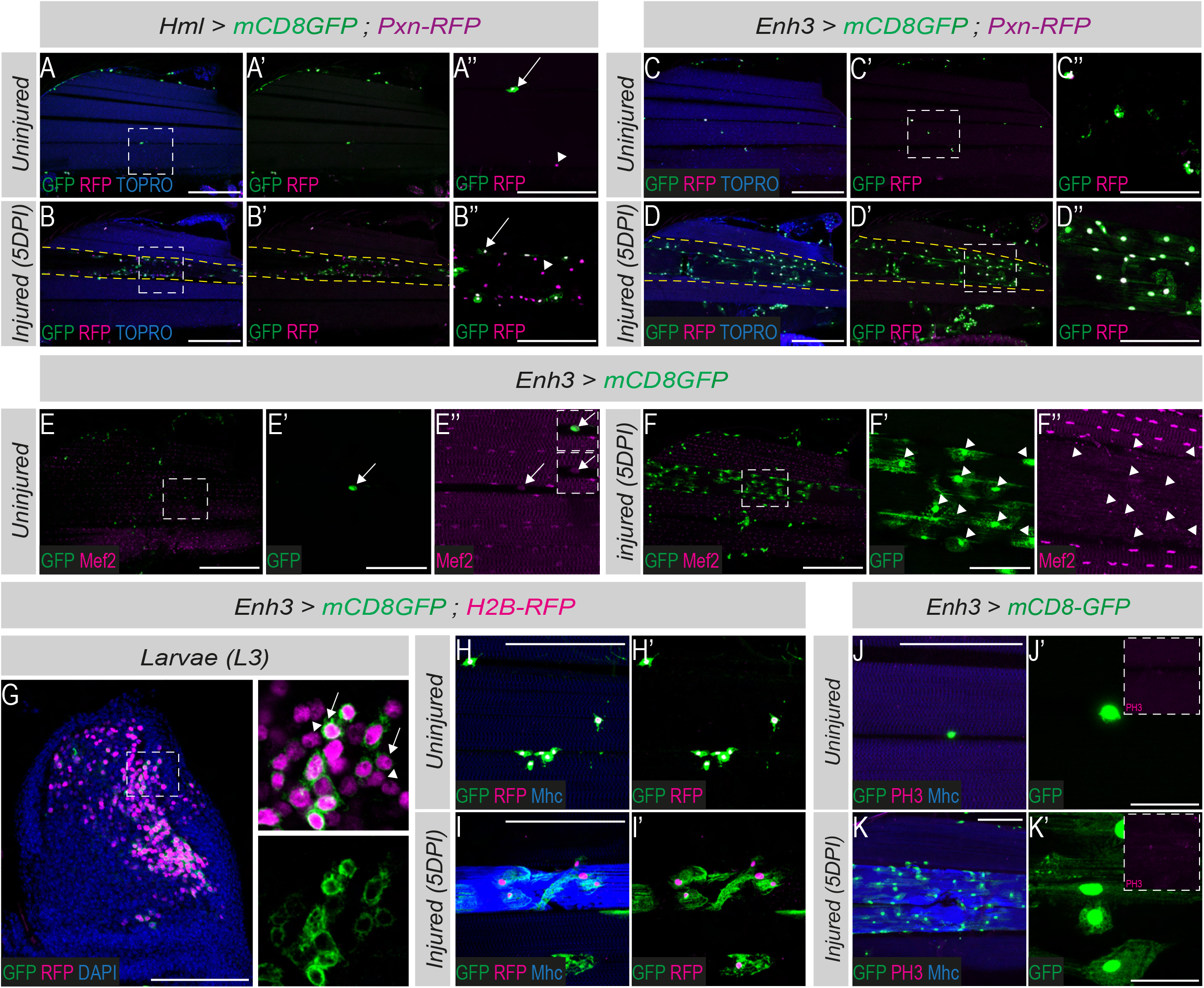
**(A-B’’).** Co-expression of hemocytes markers *HmlΔ-Gal4, UAS-mCD8GFP* (green) and *Pxn-RFP* (magenta) reveals two subsets of hemocytes associated with the IFMs: Pxn+Hml+ (arrows) and Pxn+Hml-(arrowheads). Nuclei are visualized with TOPRO (blue). Scale bars: 200 μm. (H’’-I’’). 3X magnification of the boxed regions in A and B. Scale bars: 100 μm. **(C-D’’).** Co-expression analysis of *Enh3-Gal4; UAS-mcD8GFP* (green) and *Pxn-RFP* (magenta) shows that all Enh3+ cells express Pxn. (C’’-D’’). 3X magnification of the boxed regions in C and D. Scale bars: 200 μm. **(E-E’’’).** *Enh3-Gal4; UAS-mcD8GFP* (green) expression is detected in rare adult pMPs (persistant muscle progenitors), characterized by low levels of Mef2, magenta; arrows (E’-E’’). **(F-F’’).** Recruited Enh3 positive cells (green) are negative for Mef2 expression (magneta, arrowheads). **(G).** Lineage-tracing using the ReDDM system characterization in third-instar larval wing discs (L3) (*Enh3-Gal4, UAS-mCD8GFP; UAS-H2B-RFP*). Enh3 positive cells (arrows) express both mCD8::GFP (green) and H2B::RFP (magenta), whereas Enh3 derived progeny retain only the H2B::RFP (magenta, arrowheads). Nuclei are labeled with DAPI (blue). Scale bar: 200 μm. **(H-I’).** Adult IFMs expressing the Enh3 > ReDDM lineage-tracing system under uninjured conditions (H-H’) and at 5DPI (I-I’). Muscles are stained for Mhc (blue). Recruited Enh3 potitive cells are both mCD8::GFP (green) and H2B::RFP (magenta) (n = 20 heminota from three independent replicates). **(J-K’).** Enh3 positive cells (green) are not mitotically active under either uninjured (J-J’) or injured conditions (K-K’), Anti-PH3 (magenta) is used to assess cell division and muscle fibers are visualized by Mhc staining (blue) (n = 15 heminota from two independent replicates). Scale bars: 100 μm.

**Figure S4.**
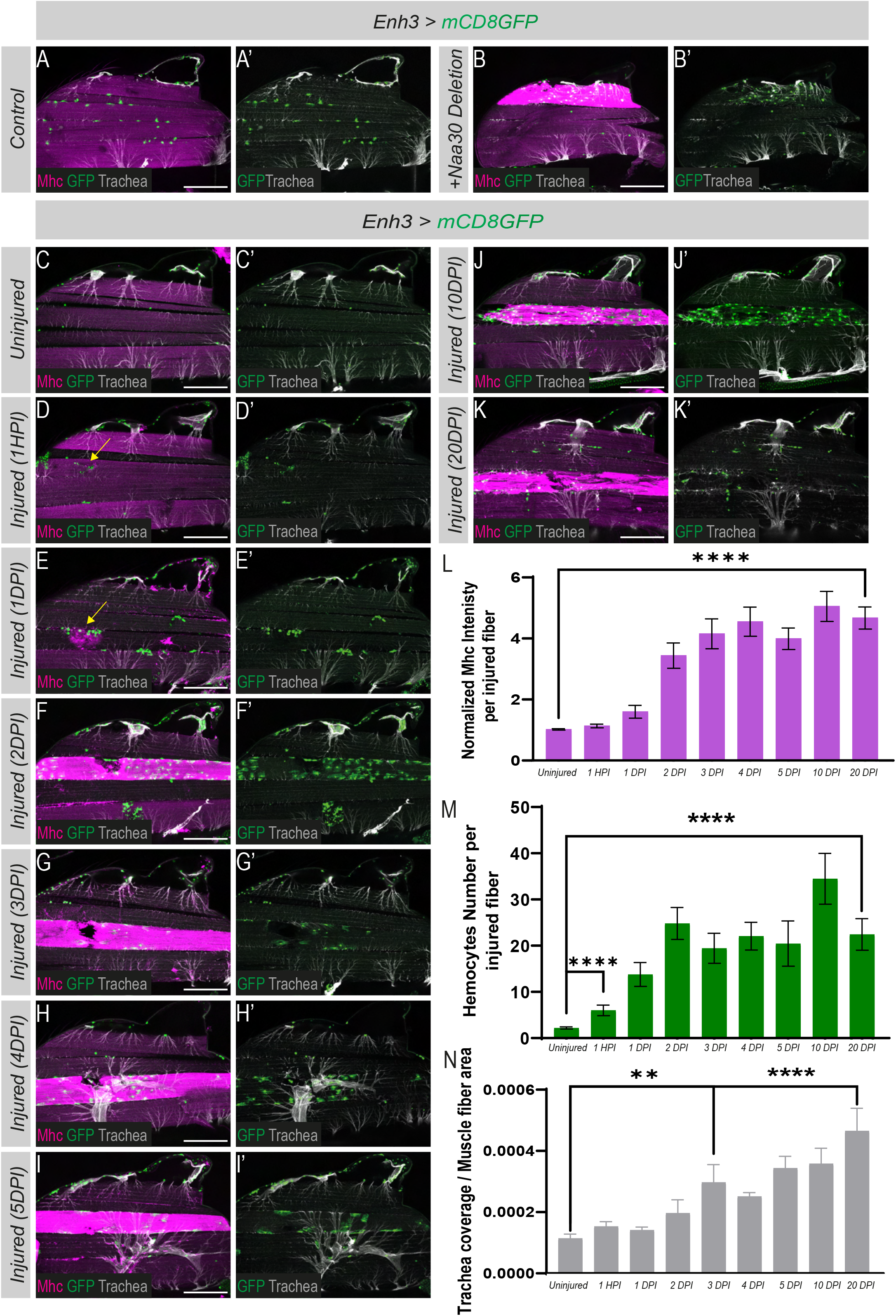
**(A-B’).** In *Naa30A mutant* flies, defective muscle attachment is associated with tracheal enrichment (gray, B’) and increased hemocytes accumulation (green, B’) compared to controls (A-A’). **(C-K’).** Time-course analysis of tracheal remodeling and hemocytes recruitment in *Enh3-Gal4; UAS-mCD8GFP* adult IFMs following localized stab injury. Tracheal branches are visualized by autofluorescence (gray), and hemocytes are labeled with GFP (green). Muscles are labeled with Mhc (magenta). Time points range from 1-hour post-injury (1HPI) to 20 days post-injury (20DPI). **(L).** Quantification of normalized Mhc intensity. **(M).** Quantification of the hemocytes number in the injured fiber shows a significant increase starting at 1HPI which remains elevated until 20DPI. **(N).** Tracheal invasion quantification shows a marked increase in its density from 3DPI and remains sustained until 20 DPI. (****p < 0.0001, ***p < 0.001; Student’s t-test; n ≥ 10 heminota per condition). Scale bars: 200 μm.

**Figure S5.**
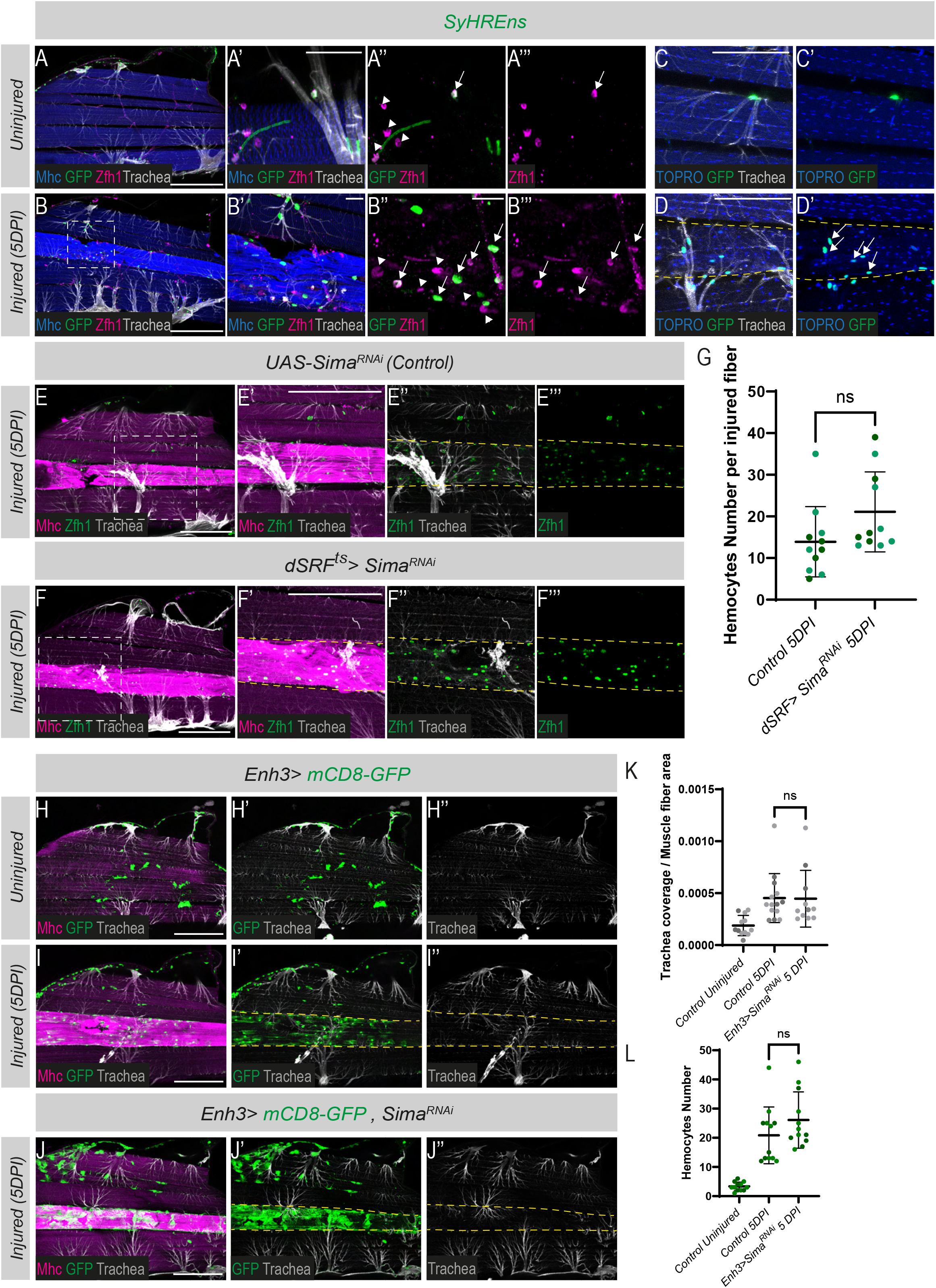
**(A–B’’’).** A subset of hemocytes marked by Zfh1 (magenta, arrows) activates the SyHREns reporter (green) in both intact and injured muscles. Arrowheads indicate hemocytes that do not activate the SyHREns reporter. Muscles are stained for Mhc (blue). Scale bars: 25 μm. **(C–D’).** SyHREns reporter activity (green) is detected in cells associated with the tracheal network but not in nuclei of injured muscle fibers (arrows). Dashed yellow outlines indicate injured muscle fibers. Scale bars: 100 μm. **(E-G).** Sima (HIF-1α) downregulation in TTCs (*dSRF-Gal4; tub-Gal80ts, UAS-Sima^RNAi^*) do not affect hemocytes (Zfh1, green) recruitment at the injured fiber. Muscle fibers are visualized by Mhc staining (magenta) and tracheal branches by autofluorescence (gray). Dashed yellow outlines indicate injured muscle fibers. Scale bars: 100 μm. **G.** Quantification of the hemocytes number (ns, Student’s t-test; n≥ 11 heminota for each condition from two independent replicates). **(H-L).** Sima (HIF-1α) downregulation in hemocytes (*Enh3-Gal4; UAS-mCD8-GFP, UAS-Sima^RNAi^*) does not affect either hemocyte recruitment nor trachea invasion into the damaged muscle fibers. Scale bars: 200 μm. **(K).** Quantification of trachea invasion into muscle fiber. **(L).** Quantification of the hemocytes number. (ns, Student’s t-test; n≥ 10 heminota for each condition from two independent replicates).

**Figure S6.**
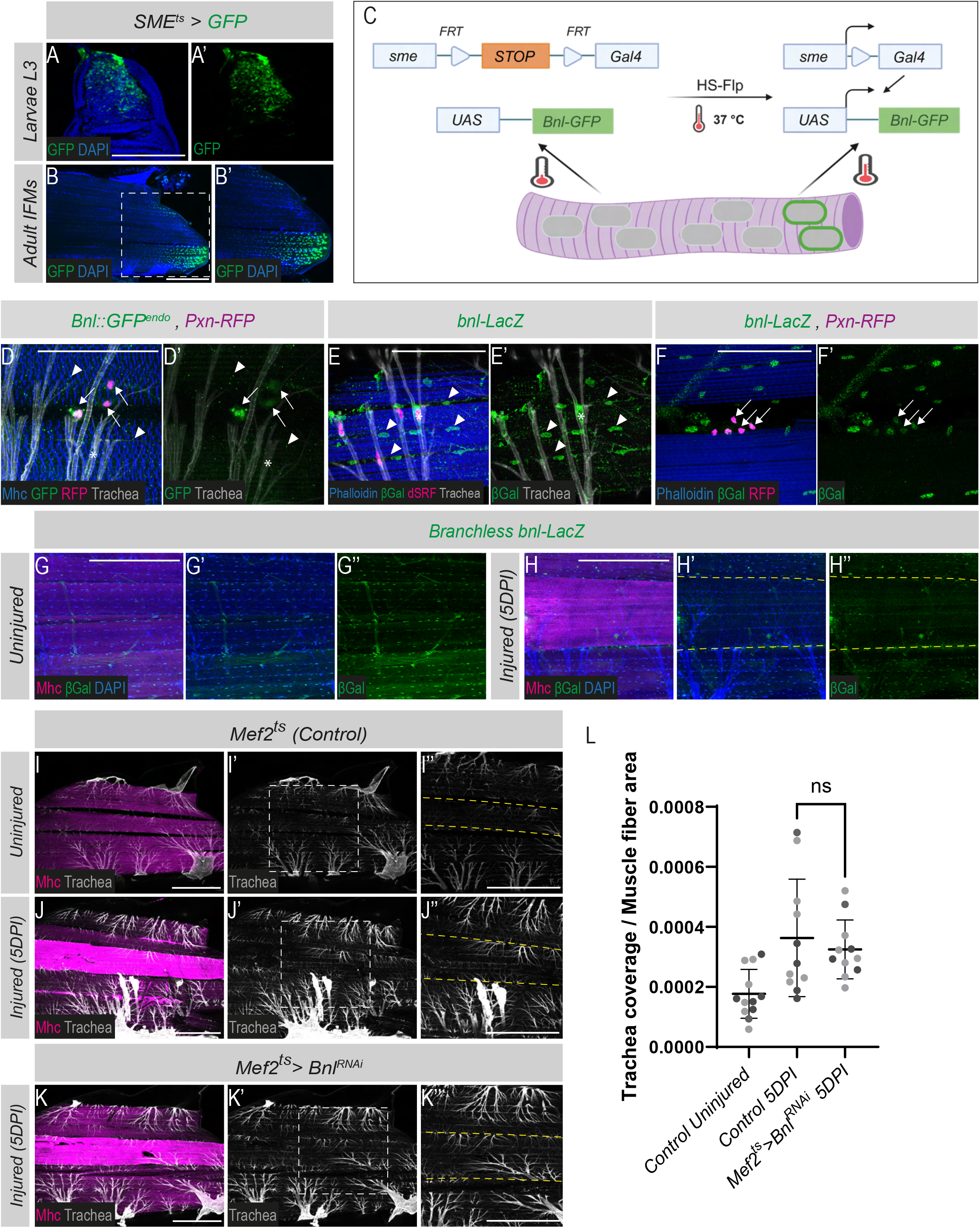
Multiple cellular sources of Bnl are present in adult IFMs. **(A-B’).** Expression pattern of the Shavenbaby (Svb) muscle regulatory enhancer (SME) (green, *SME-Gal4; UAS-mCD8GFP*). SME-Gal4 drives GFP expression (green) in a subset of larval muscle progenitors located in the posterior region of the wing disc (A-A’) and in a subset of adult muscle nuclei (green and blue (DAPI), B-B’). Scale bars: 200 μm. **(C).** Schematic representation of the FLP-OUT system used to drive UAS-Bnl::GFP expression in a subset of adult muscle nuclei, using the *Hs-FLP; FRT-SME-Stop-FRT-Gal4/CyO; tub-Gal80ts* combination. **(D-D’).** The endogenous Bnl (green, *Bnl::GFP^endo^*) is detected in IFMs, labelled with Mhc (blue; arrowheads), tracheal branches (autofluorescence, gray; asterisks), and hemocytes (*Pxn-RFP*, magenta; arrows). Scale bars: 25 μm. **(E-E’).** Expression of the *bnl-LacZ* reporter (green) is observed in terminal tracheal cells (TTCs) marked by dSRF (magenta; asterisks), and in IFMs nuclei (arrowheads). Scale bars: 25 μm. **(F-F’).** *bnl-LacZ* (green) is detected in IFMs associated hemocytes labelled with *Pxn-RFP* (magenta; arrows). Scale bars: 25 μm. **(G-H’’).** *bnl-LacZ* reporter expression (green) is downregulated in of the injured muscle fibers identified by elevated Mhc detection (magenta), the nuclei are marked with DAPI (blue). Scale bars: 100 μm. **(I-K’’).** Downregulating *bnl* in muscle (*Mef2-Gal4; tub-Gal80ts, UAS-Bnl^RNAi^*) do not alter tracheal invasion to the damaged fiber as shown in (J’’ and K’’) and quantified in **(L)** (ns, Student’s t-test; n≥ 12 heminota for each condition from two independent replicates).

**Figure S7.**
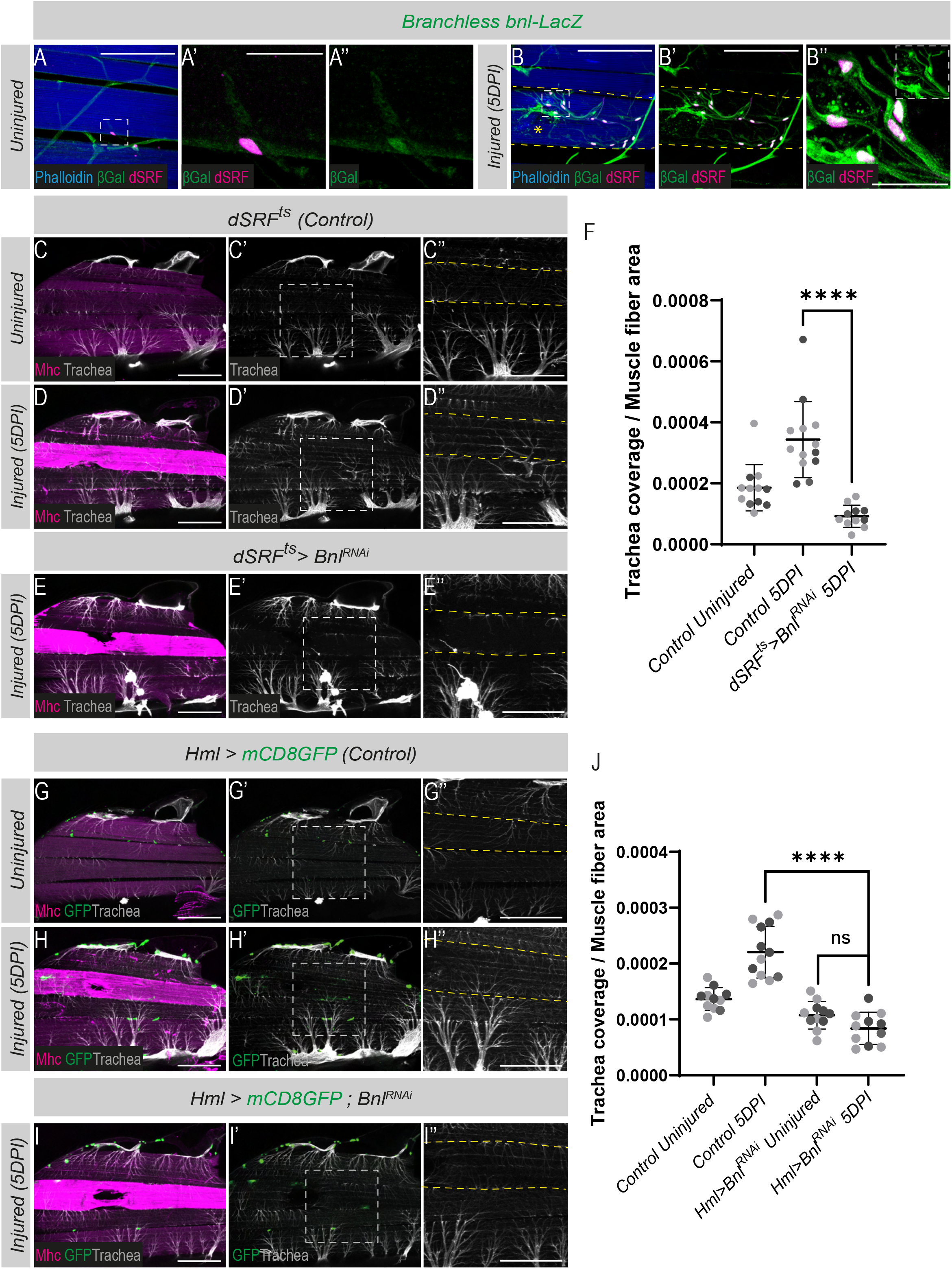
**(A–B’’).** *bnl-lacZ* reporter expression (green) is detected in tracheal terminal cells (TTCs) associated with injured muscles, identified by dSRF staining (magenta), under both control conditions and at 5 DPI. **(B’).** The tracheal branches enriched around the injured muscle fiber are positive for bnl-lacZ expression. Muscles are labelled with phalloidin (blue). Dashed yellow outlines indicate injured muscle fibers, and the asterisk marks the site of injury. **(C-F).** Downregulating *bnl* in TTCs (*dSRF-Gal4; tub-Gal80ts, UAS-Bnl^RNAi^*) decreases tracheal invasion to the damaged fiber. Muscles are labeled with Mhc (magenta), and tracheal branches are visualized via autofluorescence (gray). Scale bars: 200 μm. **(F).** Quantification of trachea invasion into the damaged fiber (****p <0.0001, Student’s t-test; n >13 heminota per condition from two independent replicates). **(G-I’’).** Targeted downregulation of *bnl* in a subset of hemocytes using *HmlΔ-Gal4* driver significantly reduces tracheal enrichment at the injured fiber, as quantified in **(J).** (****p <0.0001, Student’s t-test; n ≥ 10 heminota per condition). Scale bars: 200 μm.

**Figure S8.**
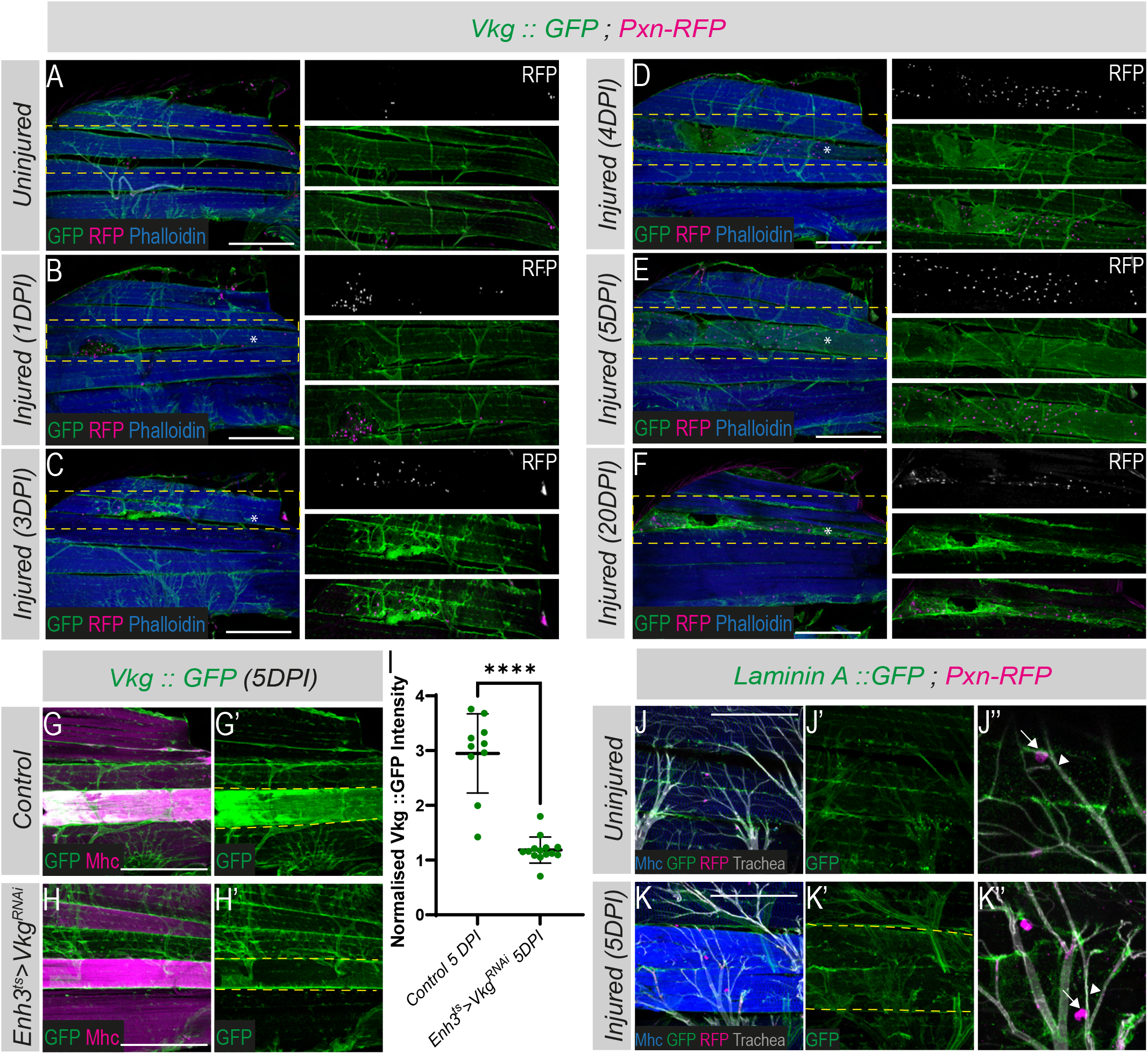
Muscle injury induces selective ECM remodelling involving Collagen IV enrichment but not Laminin A. **(A-F).** Hemocyte recruitment (magenta, Pxn-RFP) and Collagen IV distribution (green, Vkg::GFP) in uninjured muscles (A) and at successive time points following injury (B-F), from 1 to 20 DPI. Scale bar: 200 μm. **(G-G’’).** Laminin A (green, Laminin A::GFP) is detected along tracheal branches associated with IFMs (arrowheads in G’’ and H’’), but not in hemocytes (magenta, Pxn-RFP; arrows in G’’ and H’’). **(H-H’’).** At 5 DPI, Laminin A is not enriched around injured muscle fibers. Dashed yellow outlines indicate injured muscle fibers identified by elevated Mhc detection (blue). Scale bars: 200 μm. **(I-J’).** Downregulation of Collagen IV in adult hemocytes (*Enh3^ts^-Gal4 > UAS-Vkg^RNAi^*) reduces Collagen IV (green, Vkg::GFP) enrichment on the injured muscle fibers at 5DPI. Muscles are labelled with Mhc (magenta). **(K).** Quantification of normalized Vkg::GFP intensity level in injured muscle fiber at 5DPI in control flies (Vkg::GFP) and upon Collagen IV depletion (*Enh3^ts^-Gal4 > UAS-Vkg^RNAi^*) (****p<0.00001, Student t-test, n > 10 heminota per condition).

**Figure S9.**
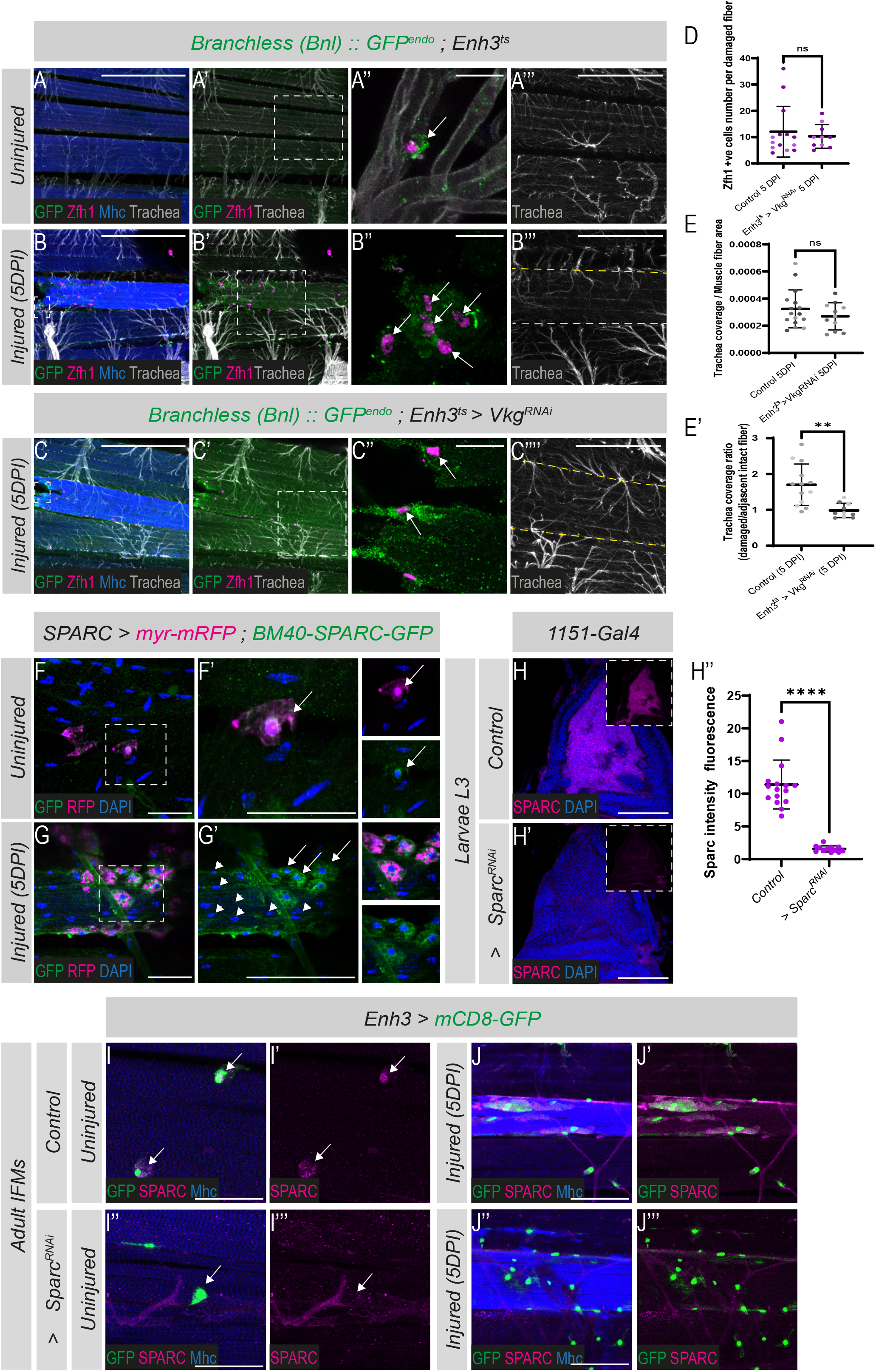
Hemocytes provide signaling and ECM components at sites of muscle injury. **(A-B’’).** Endogenous Bnl ligand distribution (green, *Bnl::GFP^endo^*) is detected around Zfh1 positive hemocytes (magenta, arrows) in intact muscles (A’’) and at 5 DPI (B’’). **(C-D).** Depletion of Collagen IV in the hemocytes using *Enh3-Gal4* driver *(Enh3-Gal4, UAS-Vkg^RNAi^)* does not affect Bnl production by hemocytes (green, Bnl::GFP^endo^; arrows) nor their recruitment to injured fibers (Zfh1, magenta) as quantified in **(D)** (ns, Student’s t-test; n ≥ 10 heminota per condition from two independent replicates). Scale bars: 200 µm. **(B’’’ and C’’’).** Hemocytes-specific knockdown of Collagen IV *(Enh3-Gal4, UAS-Vkg^RNAi^)* does not affect the trachea enrichment in the damaged fiber (Quantifications in **(E)** ns, Student’s t-test; n ≥ 10 heminota per condition from two independent replicates). The ratio of tracheal enrichment in damaged fiber relative to adjacent non-injured fiber is significantly reduced upon *vkg* knockdown **(E’)**. Scale bars: 200 µm. (**p<0.01, Student’s t-test; n ≥ 11 heminota per condition from two independent replicates). **(F-G’).** Muscle-associated hemocytes (arrows; round-shaped nuclei), but not muscle nuclei (arrowheads; flattened nuclei), express both the SPARC-Gal4, UAS-myr-RFP reporter (magenta) and BM40-SPARC-GFP (green) in uninjured and injured muscles. Nuclei are labelled with DAPI (blue). Scale bars: 25 µm. **(H-H’).** Validation of UAS-Sparc RNAi efficiency in L3 muscle progenitors (*1151-Gal4, UAS-Sparc^RNAi^*). SPARC staining intensity (magenta, H’) was significantly reduced compared with controls (H), as quantified in (H’’). (****p <0.0001, Student’s t-test; n=9). Scale bars: 100 µm. Nuclei are labelled with DAPI (blue). **(I-I’’’).** Efficient depletion of Sparc (magenta) in hemocytes (*Enh3-Gal4; UAS-mCD8GFP, green)* associated with adult IFMs. **(J-J’’’).** Hemocyte-specific Sparc depletion does not affect their recruitment to injured muscle fibers labeled by elevated Mhc staining (blue). Scale bars = 100 µm.

**Table S1. Raw tracheal coverage measurements and normalized values for all experimental conditions.**

## Reference

1. Sun F, Poss KD. Inter-organ communication during tissue regeneration. Development. 2023;150(23). Epub 20231127. doi: 10.1242/dev.202166. PubMed PMID: 38010139; PubMed Central PMCID: PMCPMC10730022.

2. Rodriguez C, Timoteo-Ferreira F, MinchioS G, Brunelli S, Guardiola O. Cellular interactions and microenvironment dynamics in skeletal muscle regeneration and disease. Front Cell Dev Biol. 2024;12:1385399. Epub 20240522. doi: 10.3389/fcell.2024.1385399. PubMed PMID: 38840849; PubMed Central PMCID: PMCPMC11150574.

3. Koike H, Manabe I, Oishi Y. Mechanisms of cooperative cell-cell interactions in skeletal muscle regeneration. Inflamm Regen. 2022;42(1):48. Epub 20221116. doi: 10.1186/s41232-022-00234-6. PubMed PMID: 36380396; PubMed Central PMCID: PMCPMC9667595.

4. Ratnayake D, Nguyen PD, Rossello FJ, Wimmer VC, Tan JL, Galvis LA, et al. Macrophages provide a transient muscle stem cell niche via NAMPT secretion. Nature. 2021;591(7849):281–7. Epub 20210210. doi: 10.1038/s41586-021-03199-7. PubMed PMID: 33568815.

5. Manneken JD, Currie PD. Macrophage-stem cell crosstalk: regulation of the stem cell niche. Development. 2023;150(8). Epub 20230427. doi: 10.1242/dev.201510. PubMed PMID: 37102706.

6. Latroche C, Gitiaux C, Chretien F, Desguerre I, Mounier R, Chazaud B. Skeletal Muscle Microvasculature: A Highly Dynamic Lifeline. Physiology (Bethesda). 2015;30(6):417–27. doi: 10.1152/physiol.00026.2015. PubMed PMID: 26525341.

7. Webster MT, Manor U, Lippincoe-Schwartz J, Fan CM. Intravital Imaging Reveals Ghost Fibers as Architectural Units Guiding Myogenic Progenitors during Regeneration. Cell Stem Cell. 2016;18(2):243–52. Epub 20151210. doi: 10.1016/j.stem.2015.11.005. PubMed PMID: 26686466; PubMed Central PMCID: PMCPMC4744135.

8. Droujinine IA, Perrimon N. Interorgan Communication Pathways in Physiology: Focus on Drosophila. Annu Rev Genet. 2016;50:539–70. Epub 20161010. doi: 10.1146/annurev-genet-121415-122024. PubMed PMID: 27732790; PubMed Central PMCID: PMCPMC5506552.

9. Boukhatmi H, Bray S. A population of adult satellite-like cells in Drosophila is maintained through a switch in RNA-isoforms. Elife. 2018;7. doi: 10.7554/eLife.35954. PubMed PMID: 29629869; PubMed Central PMCID: PMCPMC5919756.

10. Chaturvedi D, Reichert H, Gunage RD, VijayRaghavan K. Identification and functional characterization of muscle satellite cells in Drosophila. Elife. 2017;6. doi: 10.7554/eLife.30107. PubMed PMID: 29072161; PubMed Central PMCID: PMCPMC5681227.

11. Catalani E, Zecchini S, Giovarelli M, Cherubini A, Del Quondam S, BruneS K, et al. RACK1 is evolutionary conserved in satellite stem cell activation and adult skeletal muscle regeneration. Cell Death Discovery. 2022;8(1):459. Epub 20221118. doi: 10.1038/s41420-022-01250-8. PubMed PMID: 36396939; PubMed Central PMCID: PMCPMC9672362.

12. Mitchell-Gee R, Hoff R, Vishal K, Hancock D, McKitrick S, Newnes-Querejeta CV, et al. The Drosophila Him gene is essential for adult muscle function and muscle stem cell maintenance. iScience. 2026;29(2):114670. Epub 20260110. doi: 10.1016/j.isci.2026.114670. PubMed PMID: 41660269; PubMed Central PMCID: PMCPMC12874454.

13. Sanchez Bosch P, Makhijani K, Herboso L, Gold KS, Baginsky R, Woodcock KJ, et al. Adult Drosophila Lack Hematopoiesis but Rely on a Blood Cell Reservoir at the Respiratory Epithelia to Relay Infection Signals to Surrounding Tissues. Dev Cell. 2019;51(6):787–803 e5. Epub 20191114. doi: 10.1016/j.devcel.2019.10.017. PubMed PMID: 31735669; PubMed Central PMCID: PMCPMC7263735.

14. Peterson SJ, Krasnow MA. Subcellular trafficking of FGF controls tracheal invasion of Drosophila flight muscle. Cell. 2015;160(1-2):313–23. Epub 2015/01/06. doi: 10.1016/j.cell.2014.11.043. PubMed PMID: 25557078; PubMed Central PMCID: PMCPMC4577243.

15. Sauerwald J, Backer W, Matzat T, Schnorrer F, Luschnig S. Matrix metalloproteinase 1 modulates invasive behavior of tracheal branches during entry into Drosophila flight muscles. Elife. 2019;8. Epub 2019/10/03. doi: 10.7554/eLife.48857. PubMed PMID: 31577228; PubMed Central PMCID: PMCPMC6795481.

16. Hayashi S, Kondo T. Development and Function of the Drosophila Tracheal System. Genetics. 2018;209(2):367–80. doi: 10.1534/genetics.117.300167. PubMed PMID: 29844090; PubMed Central PMCID: PMCPMC5972413.

17. Affolter M, Shilo BZ. Genetic control of branching morphogenesis during Drosophila tracheal development. Curr Opin Cell Biol. 2000;12(6):731–5. doi: 10.1016/s0955-0674(00)00160-5. PubMed PMID: 11063940.

18. Perochon J, Yu Y, Aughey GN, Medina AB, Southall TD, Cordero JB. Dynamic adult tracheal plasticity drives stem cell adaptation to changes in intestinal homeostasis in Drosophila. Nat Cell Biol. 2021;23(5):485–96. Epub 2021/05/12. doi: 10.1038/s41556-021-00676-z. PubMed PMID: 33972729; PubMed Central PMCID: PMCPMC7610788.

19. Tamamouna V, Rahman MM, Petersson M, Charalambous I, Kux K, Mainor H, et al. Remodelling of oxygen-transporting tracheoles drives intestinal regeneration and tumorigenesis in Drosophila. Nat Cell Biol. 2021;23(5):497–510. Epub 20210510. doi: 10.1038/s41556-021-00674-1. PubMed PMID: 33972730; PubMed Central PMCID: PMCPMC8567841.

20. Medina AB, Perochon J, Tian Y, Johnson CT, Holcombe J, Ramesh P, et al. Neuroendocrine control of intestinal regeneration through the vascular niche in Drosophila. Dev Cell. 2025;60(22):3085–101 e6. Epub 20250721. doi: 10.1016/j.devcel.2025.06.036. PubMed PMID: 40695286.

21. Chu WC, Hayashi S. Mechano-chemical enforcement of tendon apical ECM into nano-filaments during Drosophila flight muscle development. Curr Biol. 2021;31(7):1366–78 e7. Epub 20210204. doi: 10.1016/j.cub.2021.01.010. PubMed PMID: 33545042.

22. Murphy MM, Lawson JA, Mathew SJ, Hutcheson DA, Kardon G. Satellite cells, connective Assue fibroblasts and their interactions are crucial for muscle regeneration. Development. 2011;138(17):3625–37. doi: 10.1242/dev.064162. PubMed PMID: 21828091; PubMed Central PMCID: PMCPMC3152921.

23. Orfanos Z, Sparrow JC. Myosin isoform switching during assembly of the Drosophila flight muscle thick filament lattice. J Cell Sci. 2013;126(Pt 1):139–48. Epub 20121123. doi: 10.1242/jcs.110361. PubMed PMID: 23178940.

24. Sarov M, Barz C, Jambor H, Hein MY, Schmied C, Suchold D, et al. A genome-wide resource for the analysis of protein localisation in Drosophila. Elife. 2016;5:e12068. Epub 20160220. doi: 10.7554/eLife.12068. PubMed PMID: 26896675; PubMed Central PMCID: PMCPMC4805545.

25. Varland S, Silva RD, Kjosas I, Faustino A, Bogaert A, Billmann M, et al. N-terminal acetylation shields proteins from degradation and promotes age-dependent motility and longevity. Nat Commun. 2023;14(1):6774. Epub 20231027. doi: 10.1038/s41467-023-42342-y. PubMed PMID: 37891180; PubMed Central PMCID: PMCPMC10611716.

26. Wu WH, Kuo TH, Kao CW, Girardot C, Hung SJ, Liu T, et al. Expanding the mesodermal transcriptional network by genome-wide idenAfication of Zinc finger homeodomain 1 (Zm1) targets. FEBS Lee. 2019;593(14):1698–710. Epub 20190529. doi: 10.1002/1873-3468.13443. PubMed PMID: 31093969.

27. Antonello ZA, Reiff T, Ballesta-Illan E, Dominguez M. Robust intestinal homeostasis relies on cellular plasticity in enteroblasts mediated by miR-8-Escargot switch. EMBO J. 2015;34(15):2025–41. doi: 10.15252/embj.201591517. PubMed PMID: 26077448; PubMed Central PMCID: PMCPMC4551350.

28. Ghabrial A, Luschnig S, Metzstein MM, Krasnow MA. Branching morphogenesis of the Drosophila tracheal system. Annu Rev Cell Dev Biol. 2003;19:623–47. Epub 2003/10/23. doi: 10.1146/annurev.cellbio.19.031403.160043. PubMed PMID: 14570584.

29. Jarecki J, Johnson E, Krasnow MA. Oxygen regulation of airway branching in Drosophila is mediated by branchless FGF. Cell. 1999;99(2):211–20. doi: 10.1016/s0092-8674(00)81652-9. PubMed PMID: 10535739.

30. Zhao Y, Alexandre C, Kelly G, Vincent JP, Perez-Mockus G. HIF-1alpha-mediated feedback prevents TOR signalling from depleting oxygen supply and triggering stress during normal development. Nat Commun. 2025;17(1):397. Epub 20251221. doi: 10.1038/s41467-025-67089-6. PubMed PMID: 41423448; PubMed Central PMCID: PMCPMC12796383.

31. Du L, Sohr A, Yan G, Roy S. Feedback regulation of cytoneme-mediated transport shapes a Assue-specific FGF morphogen gradient. Elife. 2018;7. Epub 20181017. doi: 10.7554/eLife.38137. PubMed PMID: 30328809; PubMed Central PMCID: PMCPMC6224196.

32. Wang X, Harris RE, Bayston LJ, Ashe HL. Type IV collagens regulate BMP signalling in Drosophila. Nature. 2008;455(7209):72–7. doi: 10.1038/nature07214. PubMed PMID: 18701888.

33. Van De Bor V, Zimniak G, Papone L, Cerezo D, Malbouyres M, Juan T, et al. Companion Blood Cells Control Ovarian Stem Cell Niche Microenvironment and Homeostasis. Cell Rep. 2015;13(3):546–60. Epub 2015/10/13. doi: 10.1016/j.celrep.2015.09.008. PubMed PMID: 26456819.

34. Gold KS, Bruckner K. Macrophages and cellular immunity in Drosophila melanogaster. Semin Immunol. 2015;27(6):357–68. Epub 20160423. doi: 10.1016/j.smim.2016.03.010. PubMed PMID: 27117654; PubMed Central PMCID: PMCPMC5012540.

35. Bunt S, Hooley C, Hu N, Scahill C, Weavers H, Skaer H. Hemocyte-secreted type IV collagen enhances BMP signaling to guide renal tubule morphogenesis in Drosophila. Dev Cell. 2010;19(2):296–306. doi: 10.1016/j.devcel.2010.07.019. PubMed PMID: 20708591; PubMed Central PMCID: PMCPMC2941037.

36. Bradshaw AD. The role of SPARC in extracellular matrix assembly. J Cell Commun Signal. 2009;3(3-4):239–46. Epub 20091002. doi: 10.1007/s12079-009-0062-6. PubMed PMID: 19798598; PubMed Central PMCID: PMCPMC2778582.

37. Chioran A, Duncan S, Catalano A, Brown TJ, Ringueee MJ. Collagen IV trafficking: The inside-out and beyond story. Dev Biol. 2017;431(2):124–33. Epub 20171002. doi: 10.1016/j.ydbio.2017.09.037. PubMed PMID: 28982537.

38. Maddaluno L, Urwyler C, Werner S. Fibroblast growth factors: key players in regeneration and Assue repair. Development. 2017;144(22):4047–60. doi: 10.1242/dev.152587. PubMed PMID: 29138288.

39. Osborn DPS, Li K, Cuey SJ, Nelson AC, Wardle FC, Hinits Y, et al. Fgf-driven Tbx protein activities directly induce myf5 and myod to initiate zebrafish myogenesis. Development. 2020;147(8). Epub 20200428. doi: 10.1242/dev.184689. PubMed PMID: 32345657; PubMed Central PMCID: PMCPMC7197714.

40. Floss T, Arnold HH, Braun T. A role for FGF-6 in skeletal muscle regeneration. Genes Dev. 1997;11(16):2040–51. doi: 10.1101/gad.11.16.2040. PubMed PMID: 9284044; PubMed Central PMCID: PMCPMC316448.

41. Lagha M, Kormish JD, Rocancourt D, Manceau M, Epstein JA, Zaret KS, et al. Pax3 regulation of FGF signaling affects the progression of embryonic progenitor cells into the myogenic program. Genes Dev. 2008;22(13):1828–37. doi: 10.1101/gad.477908. PubMed PMID: 18593883; PubMed Central PMCID: PMCPMC2492669.

42. Yablonka-Reuveni Z, Rivera AJ. Proliferative Dynamics and the Role of FGF2 During Myogenesis of Rat Satellite Cells on Isolated Fibers. Basic Appl Myol. 1997;7(3-amp4):189–202. PubMed PMID: 26052220; PubMed Central PMCID: PMCPMC4457462.

43. Zooie W, Southard SM, Braun T, Lepper C. Fibroblast growth factor 6 regulates sizing of the muscle stem cell pool. Stem Cell Reports. 2021;16(12):2913–27. Epub 20211104. doi: 10.1016/j.stemcr.2021.10.006. PubMed PMID: 34739848; PubMed Central PMCID: PMCPMC8693628.

44. Sutherland D, Samakovlis C, Krasnow MA. branchless encodes a Drosophila FGF homolog that controls tracheal cell migration and the paeern of branching. Cell. 1996;87(6):1091–101. doi: 10.1016/s0092-8674(00)81803-6. PubMed PMID: 8978613.

45. ChakrabarA S, Visweswariah SS. Intramacrophage ROS Primes the Innate Immune System via JAK/STAT and Toll Activation. Cell Rep. 2020;33(6):108368. doi: 10.1016/j.celrep.2020.108368. PubMed PMID: 33176146; PubMed Central PMCID: PMCPMC7662148.

46. Ma M, Jiang W, Zhou R. DAMPs and DAMP-sensing receptors in inflammation and diseases. Immunity. 2024;57(4):752–71. doi: 10.1016/j.immuni.2024.03.002. PubMed PMID: 38599169.

47. Evans CJ, Liu T, Girard JR, Banerjee U. Injury-induced inflammatory signaling and hematopoiesis in Drosophila. Proc Natl Acad Sci U S A. 2022;119(12):e2119109119. Epub 20220314. doi: 10.1073/pnas.2119109119. PubMed PMID: 35286208; PubMed Central PMCID: PMCPMC8944666.

48. Zhang W, Liu Y, Zhang H. Extracellular matrix: an important regulator of cell functions and skeletal muscle development. Cell Biosci. 2021;11(1):65. Epub 20210331. doi: 10.1186/s13578-021-00579-4. PubMed PMID: 33789727; PubMed Central PMCID: PMCPMC8011170.

49. Molina T, Fabre P, Dumont NA. Fibro-adipogenic progenitors in skeletal muscle homeostasis, regeneration and diseases. Open Biol. 2021;11(12):210110. Epub 20211208. doi: 10.1098/rsob.210110. PubMed PMID: 34875199; PubMed Central PMCID: PMCPMC8651418.

50. Mase A, Augsburger J, Bruckner K. Macrophages and Their Organ Locations Shape Each Other in Development and Homeostasis - A Drosophila Perspective. Front Cell Dev Biol. 2021;9:630272. Epub 20210311. doi: 10.3389/fcell.2021.630272. PubMed PMID: 33777939; PubMed Central PMCID: PMCPMC7991785.

51. Pastor-Pareja JC, Xu T. Shaping cells and organs in Drosophila by opposing roles of fat body-secreted Collagen IV and perlecan. Dev Cell. 2011;21(2):245–56. doi: 10.1016/j.devcel.2011.06.026. PubMed PMID: 21839919; PubMed Central PMCID: PMCPMC4153364.

52. Subasi BS, Grabe V, Kaltenpoth M, Rolff J, Armitage SAO. How frequently are insects wounded in the wild? A case study using Drosophila melanogaster. R Soc Open Sci. 2024;11(6):240256. Epub 20240626. doi: 10.1098/rsos.240256. PubMed PMID: 39100166; PubMed Central PMCID: PMCPMC11296199.

53. Chechenova M, McLendon L, Dallas B, Straeon H, Kiani K, Gerberich E, et al. Muscle degeneration in aging Drosophila flies: the role of mechanical stress. Skelet Muscle. 2024;14(1):20. Epub 20240820. doi: 10.1186/s13395-024-00352-4. PubMed PMID: 39164781; PubMed Central PMCID: PMCPMC11334408.

54. Siles L, Ninfali C, Cortes M, Darling DS, Postigo A. ZEB1 protects skeletal muscle from damage and is required for its regeneration. Nature Communications. 2019;10(1):1364. Epub 20190325. doi: 10.1038/s41467-019-08983-8. PubMed PMID: 30910999; PubMed Central PMCID: PMCPMC6434033.

55. Krars KP. Tissue repair: The hidden drama. Organogenesis. 2010;6(4):225–33. doi: 10.4161/org.6.4.12555. PubMed PMID: 21220961; PubMed Central PMCID: PMCPMC3055648.

56. Kirkwood TB. Understanding the odd science of aging. Cell. 2005;120(4):437–47. doi: 10.1016/j.cell.2005.01.027. PubMed PMID: 15734677.

57. Rodrigues D, Renaud Y, VijayRaghavan K, Waltzer L, Inamdar MS. Differential activation of JAK-STAT signaling reveals functional compartmentalization in Drosophila blood progenitors. Elife. 2021;10. Epub 20210217. doi: 10.7554/eLife.61409. PubMed PMID: 33594977; PubMed Central PMCID: PMCPMC7920551.

58. Lin L. Clonal Analysis of Growth Behaviors During Drosophila Larval Tracheal Development In Doctoral thesis: University of Basel; 2009.

59. Leroux E, Ammar N, Moinet S, Pecot T, Boukhatmi H. Studying Muscle Transcriptional Dynamics at Single-molecule Scales in Drosophila. J Vis Exp. 2023;(199). Epub 20230908. doi: 10.3791/65713. PubMed PMID: 37747217.

60. Chakraborty K, VijayRaghavan K, Gunage R. A Method to Injure, Dissect and Image Indirect Flight Muscle of Drosophila. Bio Protoc. 2018;8(10):e2860. Epub 20180520. doi: 10.21769/BioProtoc.2860. PubMed PMID: 34285976; PubMed Central PMCID: PMCPMC8275216.

